# Distinct disease mutations in DNMT3A result in a spectrum of behavioral, epigenetic, and transcriptional deficits

**DOI:** 10.1101/2023.02.27.530041

**Authors:** Diana C. Beard, Xiyun Zhang, Dennis Y. Wu, Jenna R. Martin, Nicole Hamagami, Raylynn G. Swift, Katherine B. McCullough, Xia Ge, Austin Bell-Hensley, Hongjun Zheng, Austin B. Lawrence, Cheryl A. Hill, Thomas Papouin, Audrey McAlinden, Joel R. Garbow, Joseph D. Dougherty, Susan E. Maloney, Harrison W. Gabel

## Abstract

Phenotypic heterogeneity is a common feature of monogenic neurodevelopmental disorders that can arise from differential severity of missense variants underlying disease, but how distinct alleles impact molecular mechanisms to drive variable disease presentation is not well understood. Here, we investigate missense mutations in the DNA methyltransferase DNMT3A associated with variable overgrowth, intellectual disability, and autism, to uncover molecular correlates of phenotypic heterogeneity in neurodevelopmental disease. We generate a DNMT3A P900L/+ mouse model mimicking a disease mutation with mild-to-moderate severity and compare phenotypic and epigenomic effects with a severe R878H mutation. We show that the P900L mutation leads to disease-relevant overgrowth, obesity, and social deficits shared across DNMT3A disorder models, while the R878H mutation causes more extensive epigenomic disruption leading to differential dysregulation of enhancers elements. We identify distinct gene sets disrupted in each mutant which may contribute to mild or severe disease, and detect shared transcriptomic disruption that likely drives common phenotypes across affected individuals. Finally, we demonstrate that core gene dysregulation detected in DNMT3A mutant mice overlaps effects in other developmental disorder models, highlighting the importance of DNMT3A-deposited methylation in neurodevelopment. Together, these findings define central drivers of DNMT3A disorders and illustrate how variable disruption of transcriptional mechanisms can drive the spectrum of phenotypes in neurodevelopmental disease.

## Introduction

As clinical sequencing becomes widely implemented and genetic studies of disease increase in scope, an expanding number of causative variants are identified in each individual disease gene. A subset of genes associated with phenotypically heterogeneous disorders such as intellectual disability and autism spectrum disorder (ASD) are primarily associated with missense mutations rather than simple truncating or loss of function mutations (Coe et al., 2019; Satterstrom et al., 2020; Wang et al., 2020). In these missense-mediated disorders, studies of multiple disease-causing alleles are important to establish phenotypes and molecular deficits that are core to all disease-associated alleles and therefore central to the disease. Isogenic animal models provide a powerful tool for understanding genotype-phenotype relationships, especially when clinical populations are small, as they minimize confounding differences from environmental factors and genetic background. Phenotypic heterogeneity and molecular differences between variants can be used to identify potential mechanisms driving the diversity of clinical presentation, while shared effects across models can define core pathways affected and provide targets for candidate therapeutic testing.

DNA methyltransferase 3A (DNMT3A)-associated neurodevelopmental disorders are an example in which it is critical to study molecular and phenotypic heterogeneity driven by a diversity of missense mutations. Heterozygous mutations within *DNMT3A* are associated with Tatton-Brown Rahman Syndrome (TBRS), an overgrowth and intellectual disability disorder typified by macrocephaly, a distinct facial gestalt, obesity, and comorbid ASD (Tatton-Brown et al., 2014, 2018). Similar to many syndromic neurodevelopmental disorder-associated genes, DNMT3A mutations have also been identified in cohorts of individuals with a primary diagnosis of ASD (Plummer et al., 2016; Sanders et al., 2012; Satterstrom et al., 2020). Disease mutations in DNMT3A are frequently missense mutations, with truncating variants or gene deletions underrepresented compared to chance estimates (Y. H. Huang et al., 2022; Tatton-Brown et al., 2018). Missense mutations generally occur within the three canonical protein domains of DNMT3A, and *in vitro* studies have demonstrated that mutations disrupt protein function through a variety of mechanisms such as altering the ability to interact with modified histones, causing loss of nuclear localization, abrogating the catalytic activity of the methyltransferase domain, or destabilizing the protein (Christian et al., 2020; Y. H. Huang et al., 2022; Lue et al., 2022). Through these mechanisms, diverse mutations may lead to differing degrees of loss of function; however, it remains unclear to what extent these mutations result in differential phenotypes *in vivo*.

Disease-associated disruption of DNMT3A likely impacts multiple aspects of nervous system development and function. DNMT3A is expressed both embryonically and postnatally and contributes to important developmental processes including genomic imprinting and maturation of the nervous system (Kaneda et al., 2004; Okano et al., 1999; Stroud et al., 2017). DNMT3A spikes in expression in postnatal neurons (between 1-3 weeks old in mice) (Feng et al., 2005; Lister et al., 2013) during which it establishes uniquely high levels of non-CpG methylation in neurons relative to other somatic cell types (Gabel et al., 2015; J. U. Guo et al., 2014; Lister et al., 2013; Nguyen et al., 2007). This methylation, predominantly at CA dinucleotides (mCA), is highly sensitive to the expression and activity levels of DNMT3A. For example, heterozygous loss of DNMT3A leads to a 50% reduction in mCA genome-wide, while modest overexpression of DNMT3A upon loss of microRNA regulation leads to excess deposition of this mark (Christian et al., 2020; Swahari et al., 2021). A major function of mCA is to recruit the methyl-binding protein MeCP2 to further regulate transcription through the activity of enhancers (Boxer et al., 2020; Clemens et al., 2019). This transcriptional regulation occurs broadly across the genome, tuning expression of large numbers of genes to allow neurons to dynamically respond to activity and maintain cell-type-specific gene expression (Gabel et al., 2015; Mo et al., 2015; Stroud et al., 2017).

Mouse models of DNMT3A disorders have established key effects associated with pathology in the human disorder, but DNMT3A missense alleles have not been systematically assessed in the brain. Analysis of DNMT3A heterozygous knockout (KO) mice has detected growth and behavioral deficits that mirror aspects of human disease, with underlying alterations in neuronal DNA methylation hypothesized to drive these effects (Christian et al., 2020; Tovy et al., 2022). In addition, recent characterization of a mouse model of the R882H mutation (DNMT3A^R878H/+^ mice) has demonstrated more severe behavioral disruption in comparison to the heterozygous KO (Christian et al., 2020; Smith et al., 2021). However, studies of the R882H mutation in acute myeloid leukemia suggest this mutation results in dominant negative effects not observed for other mutations (Emperle et al., 2018; Russler-Germain et al., 2014; Smith et al., 2021). Furthermore, the impact of the R882H mutation on neuronal DNA methylation has not been assessed, and no model representing the majority of “typical” missense mutations causing partial loss of function has been systematically analyzed *in vivo*. Therefore, core deficits shared by the majority of disease-associated DNMT3A missense mutations remain undefined, and the molecular underpinnings driving a spectrum of severity have not been assessed.

In this study, we addressed these gaps in knowledge by generating and characterizing a mouse model of DNMT3A P904L mutation. Using these mice, which model a class of missense mutations partially disrupting the methyltransferase activity of DNMT3A (Christian et al., 2020; Tatton-Brown et al., 2018), we defined core deficits that are observed across DNMT3A models, including overgrowth, obesity, social alterations, and reductions of neuronal DNA methylation. We compared this model to a model of R882H mutation to identify distinct phenotypic, epigenomic, and transcriptional effects of these two DNMT3A missense mutations *in vivo* and observed more dramatic impacts on enhancer activity linked to increased phenotype severity in the R882H model. Finally, we leverage these datasets to establish common molecular pathways and etiology shared within DNMT3A models and use these core effects to explore convergent molecular mechanisms potentially contributing to nervous system disruption across related neurodevelopmental disorders.

## Results

### P900L heterozygous mutant mice exhibit obesity and bone length overgrowth consistent with other DNMT3A models

To investigate the range of effects caused by missense mutations in DNMT3A and characterize a “typical” mutation, we used CRISPR/Cas9 technology to generate a constitutive DNMT3A P900L/+ (P900L) mutant mouse mimicking the recurrent human P904L mutation (see methods). Sanger sequencing confirmed the correct heterozygous mutation (Supplemental Figure 1A-B). This mutation has been shown to have robust loss of function effects when characterized in vitro (Christian et al., 2020), and P900L mutants did not display severe changes in general health (Supplemental Figure 1C). RT-qPCR of transcript levels of DNMT3A P900L mice showed no detectable differences, however a subtle reduction in protein expression was observed by western blot (Supplemental Figure 1D). This mild reduction in protein expression indicates that though the mutant protein is expressed, a subset of effects may be caused by this reduction in overall DNMT3A levels. With this model in hand, we next assessed phenotypic differences caused by the P900L mutation.

Patients with DNMT3A mutations exhibit overgrowth (defined as being +2 standard deviations from mean height), macrocephaly, and a distinctive facial gestalt, therefore we measured homologous morphological changes in the P900L model to determine if these mutants displayed similar overgrowth phenotypes. Human height is well correlated with leg bone length (Duyar & Pelin, 2003), therefore we used digital x-ray imaging to measure femur length and found that P900L femurs were significantly longer than WT littermates (Figure 1A-B). We examined changes in skull morphology for macrocephaly using µCT imaging and found no overall increase in skull size, with few very subtle changes in distances between Euclidian landmarks on the skull (Figure 1C-D). This indicates that the P900L model exhibits significant changes in long bone length, without overall changes in skull size or shape.

**Figure 1:**
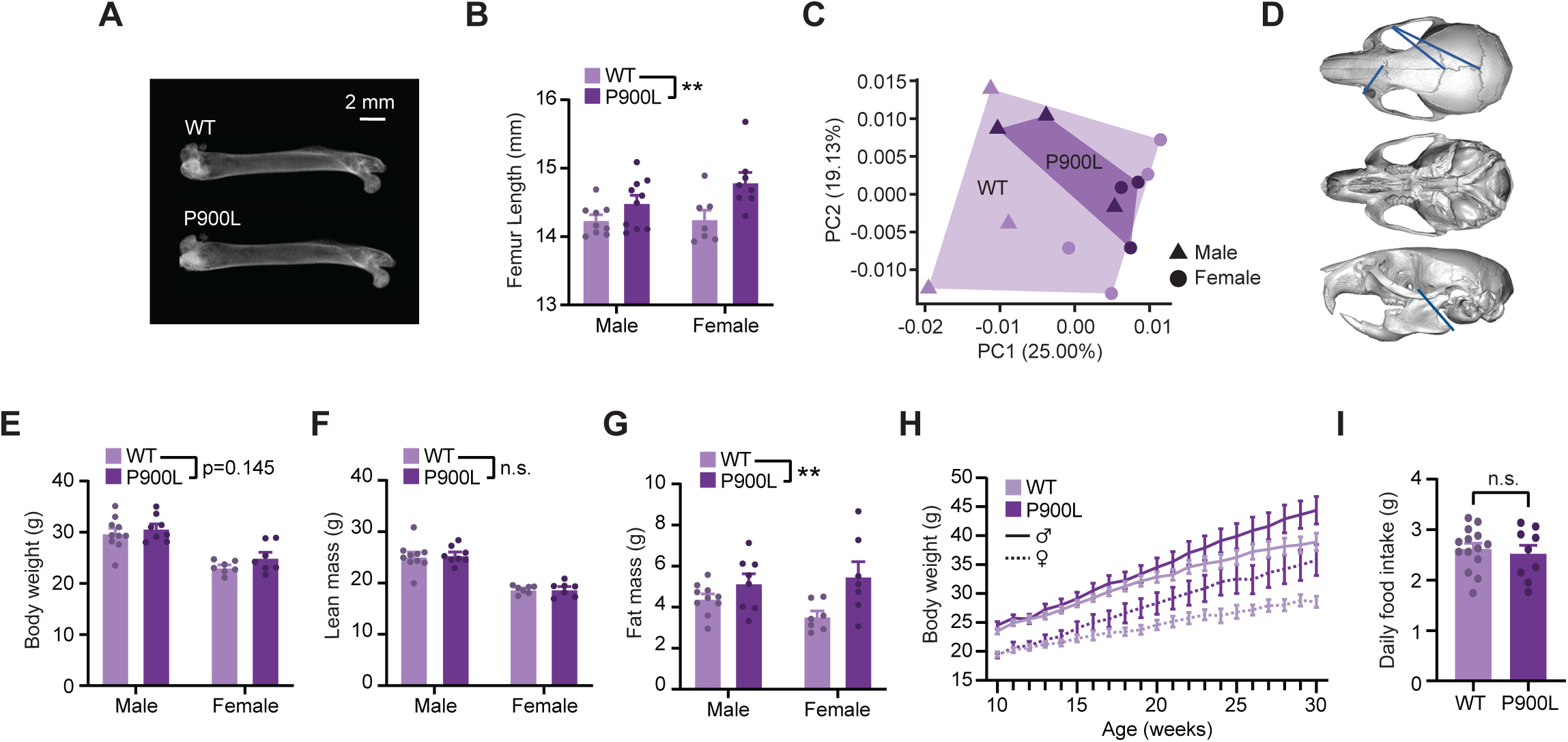
P900L mutants have increases in bone length and progressive obesity. (A) Representative dual x-ray image of femurs isolated from 30-week WT and P900L littermates. (B) Quantification of femur length (P900L n=18, 10 male, 8 female; WT n=16, 9 male, 7 female; 2-way ANOVA, genotype ** *p*<0.01). (C) Principal components analysis of skull landmark distances (P900L n=8, 4 male, 4 female; WT n=8, 4 male, 4 female). (D) Example image of reconstructed skull from µCT imaging with significant linear distances shown. Blue lines indicate distances that were significantly longer in the WT compared to the P900L, and no distances were significantly longer in the P900L (E-G) Quantification of body weight (E) and EchoMRI measures of lean mass (F) and fat mass (G) (P900L n=15, 8 male, 7 female; WT n=17, 10 male, 7 female; 2-way ANOVA, genotype ** *p*<0.01). (H) Body weights of animals on a high-fat diet measured weekly for 20 weeks (P900L n=18, 10 male, 8 female; WT n=16, 9 male, 7 female; 3-way repeated measures ANOVA; genotype *p*=0.022; genotype by time *p*<0.0001) (I) Daily food intake between 30-week WT and P900L animals that had been on a high fat diet for 20 weeks is not significantly changed (P900L n=9, 3 male, 6 female; WT n=14, 7 male, 7 female; Unpaired T-Test *p*=0.624). Results are expressed as mean ± SEM. No significant sex-genotype interactions observed.

Obesity is an emerging phenotype in the clinical TBRS population, and it has the potential to impact many aspects of health for patients and families. To examine if an obesity phenotype exists in the P900L model, we measured body weight and used EchoMRI to measure body mass content in adult animals. P900L mutants displayed trends towards increased body weight at 30-35 weeks and a significant increase in fat mass with no change in lean mass, indicating that these animals have an obesity phenotype (Figure 1E-G). High fat diets can exacerbate progressive weight gain effects (Smith et al., 2021) therefore we next measured weight gain over time in animals on a high fat diet and found that there was a significant increase in weight gain in the P900L animals compared to their WT littermates (Figure 1H). Notably, this increase in body weight does not appear to be driven by an increase in food consumption (Figure 1I). Thus, this new mutant model exhibits a progressive obesity phenotype that may be driven by metabolic or cellular changes, rather than a difference in feeding behavior.

These findings indicate that the P900L mutant has increases in long bone length and body fat, suggesting that DNMT3A-associated overgrowth and obesity is consistent across multiple mutations and can be studied using mouse models (Christian et al., 2020; Smith et al., 2021; Tovy, Reyes, et al., 2022). In further agreement with published models, the P900L has no dramatic differences in skull size or shape, suggesting that DNMT3A mouse models may not be ideal for investigating TBRS-associated cranial overgrowth or facial structure. In contrast, the existence of a reproducible increase in long bone length and body fat across this and other mouse models indicates conserved DNMT3A-dependent processes affecting body fat and skeletal development.

### DNMT3A mutant mice have decreased brain volume

Macrocephaly is a common phenotype in TBRS, and other structural brain changes such as ventriculomegaly and Chiari malformation have been observed (Tatton-Brown et al., 2014, 2018); however, brain size and structure in mice have yet to be investigated. We therefore interrogated whether brain size or structure are affected in P900L adult mice using anatomic magnetic resonance imaging (MRI) and diffusion tensor imaging (DTI). No gross structural changes were detected, and surprisingly, P900L mutant mice exhibited decreased whole brain volumes (Figure 2A-B). We also observed trends towards reduced volume of the corpus callosum that were proportional to changes in whole brain volume (Figure 2C-D) and found a subtle but significant decrease in mean corpus callosum fractional anisotropy (FA) in the P900L mutant (Figure 2E) indicating potential changes in white-matter tract integrity or organization. P900L mutants also exhibited reduced cortical thickness across multiple cortical regions (Figure 2F-G). No significant differences in ventricular volume were detected (Figure 2H-I). Together this phenotype contrasts with clinical data but suggests fundamental developmental processes impacting brain size are disrupted by DNMT3A mutation in mice.

**Figure 2:**
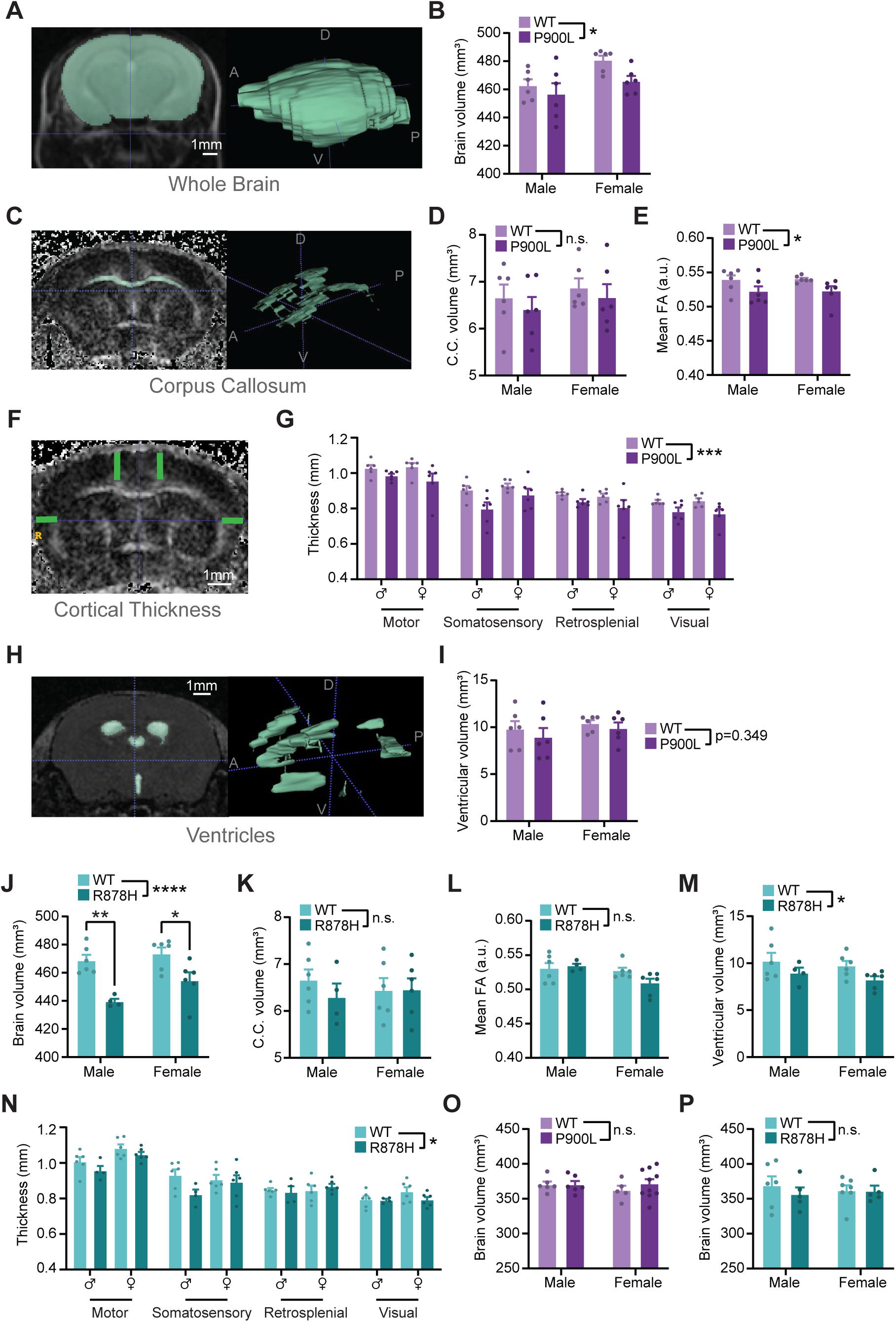
DNMT3A mutations cause reductions in brain volume and cortical thickness. (A) Representative MRI image (left) and whole brain segmentation (right). D-Dorsal, V-Ventral, A-Anterior, P-Posterior (B) Quantification of whole brain volume from WT and P900L adults. (C) Representative MRI image of fractional anisotropy (FA) of corpus callosum (left) and example segmentation (right). (D-E) Quantification of corpus callosum volume (D) and FA (E) in WT and P900L animals. (F) Representative image of cortical thickness measurements. (G) Quantification of cortical thickness across various regions (3-way repeated measures ANOVA, genotype *p*<0.0001, region *p*<0.0001). (H) Representative image of ventricles from an ADC image (left), and example segmentation (right) (I) Quantification of ventricular volume in WT and P900L animals (J-N) Quantification of WT and R878H whole brain volume (J), corpus callosum volume (K) and FA (L), ventricular volume (M) and cortical thickness (N) (Cortical thickness: 3-way repeated-measures ANOVA, genotype *p*<0.05, region *p*<0.0001) (O-P) Brain volume measurements at P10 shows no significant difference between P900L mutants (O) or R878H mutants (P) and their WT littermates. Results are expressed as mean ± SEM with individual animals shown. Genotype effect from 2-way ANOVA and significant within-sex comparisons with Sidak’s multiple testing correction are shown. As some measures showed a significant sex-effect, sexes were separated, and sex is included as a factor. Detailed statistics, and sample sizes in Supplemental Table 1. * *p*<0.05; ** *p*<0.01; *** *p*<0.001; **** *p*<0.0001

Because brain volume has not been previously investigated in DNMT3A mutant mice, we repeated MRI imaging and volumetric analyses in DNMT3A^R878H/+^ (R878H) mice mimicking the severe R882H mutation (Smith et al., 2021). Consistent with the P900L mutant, R878H mutants showed a significant reduction in brain volume (Figure 2J), though there were no significant changes in corpus callosum size or FA (Figure 2K-L) suggesting that white-matter effects observed in the P900L mutants may be allele-specific. However, R878H mutants did have significantly reduced ventricular volume and cortical thickness (Figure 2M-N), which likely contributes to the overall reduction in brain volume. These data demonstrate that reduced brain volume is a shared phenotype in DNMT3A mutant mice.

Given that DNMT3A has critical roles both embryonically (Kaneda et al., 2004; Okano et al., 1999) and in early postnatal development (Lavery et al., 2020; Stroud et al., 2017), we next assessed if onset of brain volume phenotypes occurs during early development or arises progressively. This is especially important because previous studies have demonstrated transcriptional overlap between DNMT3A and MeCP2 disorders, and MeCP2 mutants have progressive decrease in brain volume which may be phenocopied in DNMT3A mutants (Akaba et al., 2022; Allemang-Grand et al., 2017; Christian et al., 2020). We therefore imaged DNMT3A mutant mice early in postnatal development, after any embryonic actions of DNMT3A and before postnatal DNMT3A or MeCP2 are highly expressed. MRI analysis at P10 found no evidence of altered brain size in P900L or R878H mutants (Figure 2O-P). Because the disruption of brain volume occurs between P10 and adulthood in DNMT3A mutants, these results suggest that reduced brain volume may be due to deficits in postnatal maturation or survival rather than generation of neural cells.

### P900L mutants have altered social behavior and tactile sensitivity without changes in activity, anxiety-like behaviors, or learning and memory

Like numerous other syndromes associated with intellectual disability and ASD, DNMT3A disorders are typified by variable behavioral deficits and there is limited understanding of the molecular drivers of this diversity. To begin to define the shared and distinct cognitive phenotypes, we first focused on domains of behavior previously disrupted in other models of DNMT3A disorders, including activity, exploration, and anxiety-like behaviors (Christian et al., 2020; Nguyen et al., 2007; Smith et al., 2021). We measured activity and natural digging behaviors using an open field assay and marble burying assay, respectively. We found that P900L mutants traveled similar distances compared to WT littermates (Figure 3A) and had no differences in digging behavior (Figure 3B). No motor phenotypes were observed using continuous or accelerating rotarod assays (Supplemental Figure 2A-B), and mutant mice showed no differences in coordination or broad sensorimotor measures (Supplemental Figure 2C-H). To assess anxiety-like behaviors, we measured the time in the center of an open field and the time in open arms of an elevated plus maze and found that P900L mutants showed no significant differences in either of these assays (Figure 3C-D). These results indicate that exploration, motor, and anxiety phenotypes are not shared across all DNMT3A models.

**Figure 3:**
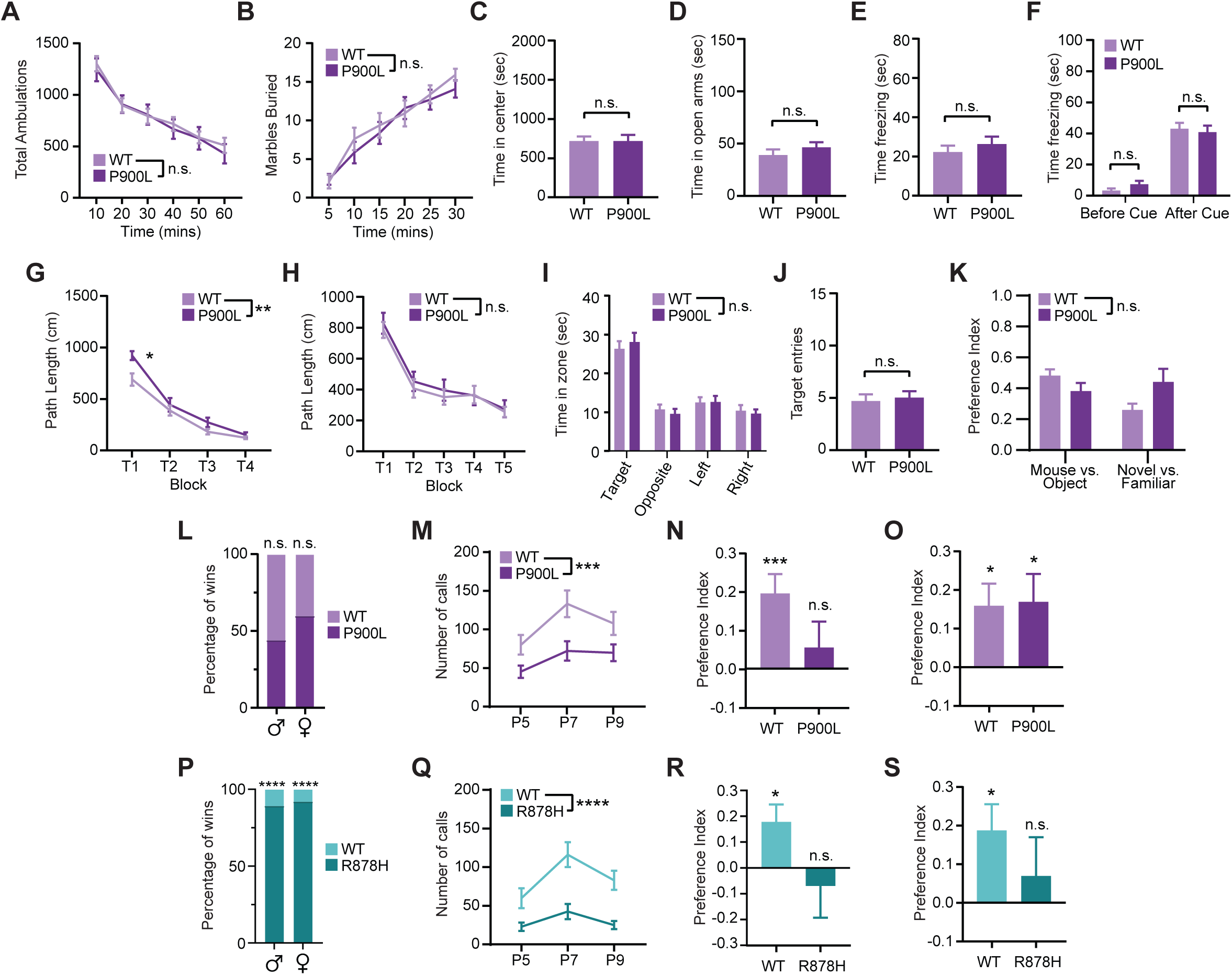
P900L mutants do not show activity or anxiety-like phenotypes, but do have changes in social and tactile behavior. (A) Measurement of movement in an open field assay over 60 minutes. (B) Marble burying behavior over a 30-minute period. (C) Time in the center of an open field assay. (D) Time in the open arms of an elevated plus maze. (E-F) Time spent freezing in a conditioned fear assay. Animals were trained to associate an environment and a stimulus (tone) with a small foot-shock, and freezing behavior was recorded. P900L mutants had similar time spent freezing compared to WT littermates for both the environmental context alone (E), and in response to the tone (cue) stimuli (F). (G-J) Measures of spatial learning and memory in a Morris Water Maze. During cued trials (G) the escape platform was visible, and path length to escape platform was measured. P900L animals had significantly longer path length when initially exposed to the task, but no differences by trial 4. During place trials (H) the platform was no longer visible, and genotypes had similar path lengths to reach the escape platform. (I) Time in quadrants after the platform was removed indicates no significant differences in target zone, and (J) both genotypes crossed over where the escape platform had been located a similar number of times. (K) In a 3-chamber social approach assay, P900L and WT animals both had a similar preference index for a mouse rather than an object, and for a novel mouse rather than a familiar one. Both genotypes showed a significant non-zero preference for interacting with conspecific mouse over an object, and for interacting with a novel conspecific over a familiar conspecific. All preference indexes range from −1 to 1. (L) Tube test percentage of bouts won, indicating that WT and P900L animals were equally likely to win bouts. (n=140 bouts) (M) Ultrasonic vocalizations of P900L and WT P5-P9 pups when isolated from the nest. (N) Preference index for the novel object during a tactile novel object recognition assay for WT and P900L animals. One-sample T-Test to determine if preference index is significantly different than 0 is indicated for both genotypes. (O) Preference index for a visually distinct novel object for P900L and WT animals. One sample T-Test to determine if preference index is significantly different than 0 is indicated. (P) Tube test percentage of bouts won, indicating that R878H animals won significantly more bouts than WT animals. (n=51 bouts) (Q) Ultrasonic vocalizations of WT and R878H pups when isolated from the nest. (R) Preference index for the novel object during a tactile novel object recognition task for WT and R878H animals. One-sample T-Test to determine if preference index is significantly different than 0 is indicated for both genotypes. (S) Preference index for visually distinct novel object for R878H and WT animals. One sample T-Test to determine if preference index is significantly different than 0 is indicated Graphs indicate mean ± SEM. Detailed statistics, and sample sizes in Supplemental Table 1. * *p*<0.05; ** *p*<0.01; *** *p*<0.001; **** *p*<0.0001

Intellectual disability is a central phenotype used for clinical diagnosis of TBRS that may be variably present in individuals identified with DNMT3A mutations through studies of ASD. We next examined whether the P900L mutant displayed deficits in the conditioned fear and Morris water maze assays to assess aversive associative memory and spatial learning and memory, respectively. In the conditioned fear assay, the P900L mutants displayed normal responses to aversive stimulus presentation, and normal contextual and cued fear memory (Figure 3E-F; Supplemental Figure 2I). P900L mutants also showed normal spatial learning in the Morris water maze assay, as exhibited by mutants learning the location of a visually cued platform and recalling the location of a hidden platform, following a slight difference upon initial task exposure (Figure 3G-H). When the hidden platform was removed, mutants and WT mice spent similar times investigating each quadrant of the maze with similar number of entries into the zone where the platform was previously located (Figure 3I-J). Notably, the absence of robust learning and memory deficits in the P900L mutation appears to mirror individuals with the homologous human P904L mutation that do not have intellectual disability diagnoses (Sanders et al., 2015).

Because mutations in DNMT3A are not only associated with intellectual disability but are also identified in individuals with a primary diagnosis of ASD (Sanders et al., 2012; Satterstrom et al., 2020), we next assessed common phenotypes displayed in mice carrying mutations in autism-associated genes (Chen et al., 2021; Han et al., 2020). In a three-chamber social approach assay, the P900L mutants and WT littermates showed similar preferences for exploring a conspecific rather than an object, and for exploring a novel conspecific rather than a familiar mouse, with no change in overall distance traveled (Figure 3K; Supplemental Figure 2J) suggesting that mutants have no change in social preference. P900L and WT mice both won a similar number of bouts in the tube test, indicating no broad changes to social dominance or hierarchies (Figure 3L; Supplemental Figure 2K). However, when we measured isolation-induced vocalizations in mouse pups, we found that mutant pups call significantly less when removed from the nest indicating deficits in early communication behaviors (Figure 3M). Overall spectral, temporal, and body weight features were largely unchanged, with slight decreases in volume, suggesting no major motor, developmental, or respiratory deficits are driving this social phenotype (Supplemental Figure 2L-N). These results show that the P900L mutation causes significant deficits in neonatal communication behavior.

Effects in somatosensory processing have been implicated as a driver of behavioral phenotypes in ASD, and mice carrying mutations in ASD-associated genes have been shown to have deficits in tactile discrimination (Orefice et al., 2019; Orefice et al., 2016). Therefore, we next measured tactile discrimination using a textured novel object recognition (NORT) task in which mice explore objects that are visually indistinguishable but differ in texture. We found that WT mice showed a preference to explore a novel tactile object, and this preference was lost in the P900L mutant (Figure 3N; Supplemental Figure 2O). To directly test if this lack of preference was due to differences in tactile discrimination or more general novelty discrimination deficits, we re-ran this task using visually and physically distinct objects and found that mutant and WT littermates displayed similar preferences for novel objects (Figure 3O; Supplemental Figure 2P). This indicates that P900L mutants have alterations in tactile discrimination rather than broad associative memory deficits or changes in novelty-seeking behaviors. This specific deficit in somatosensory processing indicates potential changes in the peripheral nervous system or sensory processing and suggests such deficits may contribute to autism phenotypes in affected individuals.

The R878H mutation generally causes more severe behavioral deficits than those observed here in the P900L mutant (Smith et al., 2021); however, it remains unknown if the R878H mutant demonstrates social or tactile phenotypes. Therefore, we tested the R878H mutant model for changes in ultrasonic vocalizations, social hierarchies, and tactile discrimination. We found that this model exhibited robust changes in social hierarchies, as shown by mutants overwhelmingly winning bouts in the tube test (Figure 3P). Interestingly, this change in behavior is not driven by a change in body weight, as DNMT3A mutants develop late onset obesity and did not have a significant difference in body weight at the time of testing (Supplemental Figure 2Q). Furthermore, in the majority of R878H wins, the mutant mouse actively moved into the tube and pushed the other mouse out, indicating that the wins were not due to lack of movement or activity (Supplemental Figure 2R). To further examine social phenotypes, we next measured pup ultrasonic vocalizations and found that R878H mutants also called significantly less than WT littermates when isolated from the nest, indicating decreases in pup communication (Figure 3Q). Pups showed no major deficits in motor or respiratory measures, with slight decreases in mean frequency of calls, indicating that major motor or respiratory phenotypes are not driving this decrease in communication (Supplemental Figure 2S-U). However, the R878H mutants did weigh significantly less than WT littermates during this developmental window (Supplemental Table 1), thus we cannot rule out developmental delay as an underlying cause of this phenotype. Finally, we assessed tactile discrimination and found that R878H mutant mice, unlike WT littermates, have no preference for a novel tactile object (Figure 3R; Supplemental Figure 2V). R878H mutants also lack a preference for visually distinct novel objects, indicating that this may potentially be a broader associative learning and memory disruption (Figure 3S; Supplemental Figure 2W). These data demonstrate that the R878H and P900L mutants share communication and tactile discrimination deficits.

Together, our findings indicate that the P900L mutation does not cause deficits in activity, exploration, or anxiety-like behaviors, contrasting previous findings for R878H mutants (Smith et al., 2021) and DNMT3A heterozygous KO mice (Christian et al., 2020), suggesting that these are not central phenotypes consistently associated with DNMT3A disruption. We also identify robust changes in social hierarchies in the R878H mutant, which in combination with previous work, supports uniquely strong phenotypes in this mutant. These results suggest that the R878H and KO mutations are more severe than the P900L, but all mutants display some disease-relevant behavioral deficits such as decreases in ultrasonic vocalizations and loss of preference for novel tactile objects. Additionally, shared phenotypes may be particularly important measures for future work investigating cellular and molecular changes that may be contributing to disease.

### Differential alterations in DNA methylation mirror differential phenotypic severity in DNMT3A mutant mice

Given the critical role DNA methylation plays in nervous system function, we hypothesized that altered methylation in the brain is a central driver of disease, and that differential levels of disruption between mutants may contribute to variable phenotypic severity. We therefore examined how P900L and R878H mutations affect DNA methylation in the brain by using whole genome bisulfite sequencing to measure DNA methylation across a variety of brain regions (Figure 4A-B). Strikingly, in all brain regions the P900L mutants had a ∼50% reduction of genome-wide mCA, and the R878H mutants had an even more severe ∼75% reduction of CA methylation (Figure 4A). Effects on global mCG levels were less dramatic, with trending reductions of global mCG in the P900L mutant and significant reductions in the R878H mutant (Figure 4B). Thus, mutation of DNMT3A causes a widespread reduction of neuronal methylation, and mCA levels are particularly sensitive to DNMT3A disruption.

**Figure 4:**
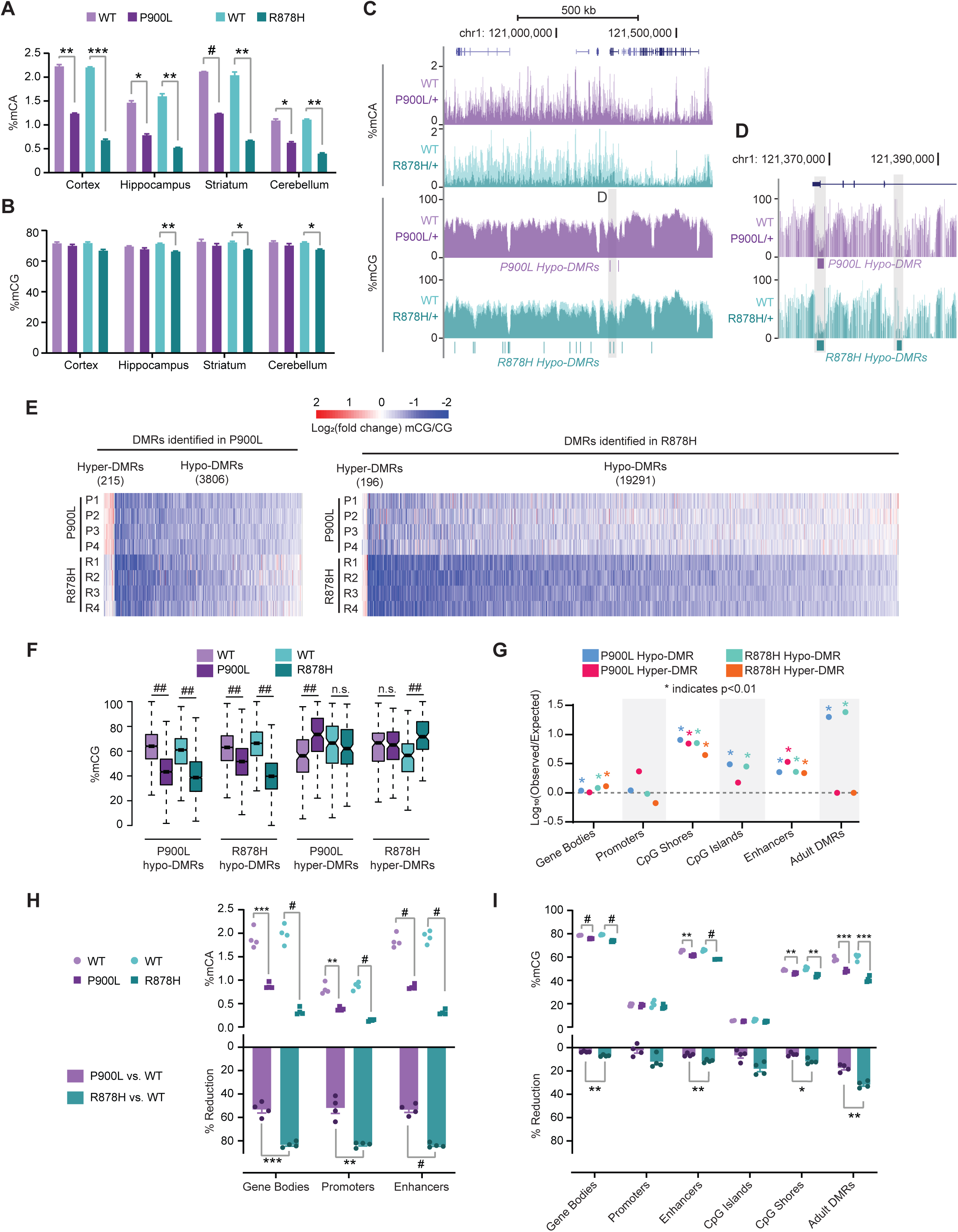
DNMT3A mutants have significant changes to DNA methylation, with more extreme changes in the R878H mutant compared to the P900L. (A-B) Average genome-wide methylation levels from brain regions measured using whole genome bisulfite sequencing for (A) percent mCA and (B) percent mCG for both P900L and R878H mutants and their WT littermates. (All groups n=4, 2 male, 2 female; Unpaired Student’s T-Test with Bonferroni Correction) (C-D) Representative genome browser view showing percent mCA and mCG (C). (D) Zoomed in browser to show changes in CG at hypo-differentially methylated regions (DMRs). (E) Heatmap of CG-DMRs identified in P900L and R878H mutants vs. their WT littermates. Log_2_(Fold change mCG/CG) indicated between each littermate pair for each DMR. (F) Average mCG in each genotype at DMRs called in both mutant strains. Both mutants show a consistent decrease at hypo-DMRs called in either mutant. Hyper-DMRs are only significant in the strain they were defined in. (G) Overlap analysis of DMRs with genomic regions of interest. Adult DMRs are the regions that significantly gain mCG over postnatal development in neurons (Lister et al. 2013). No point indicated for R878H Hyper-DMRs CpG islands, due to 0 resampled DMRs overlapping. Significance assessed with a Chi-Squared test with expected proportions of overlapping and nonoverlapping measured by resampling DMRs (H) Average mCA level at regions of interest (top) and percent reduction of mCA between WT and mutants (bottom). (I) Average mCG level at regions of interest (top) and percent reduction of mCG between WT and mutants (bottom). Promoters and CpG islands have low levels of mCG, and a trending, but not significant difference between P900L and R878H loss. Bar graphs indicate mean ± SEM. Notched box and whisker plots indicate median, interquartile, and confidence interval of median. All groups n=4, 2 male, 2 female. Detailed statistics, and sample sizes in Supplemental Table 1. * *p*<0.05; ** *p*<0.01; *** *p*<0.001; # *p*<0.0001; ## *p*<2 e^-10^

To systematically assess altered DNA methylation and its potential impact on gene regulation we next interrogated methylation changes at kilobase scale regions including enhancers and gene bodies. We performed high-depth sequencing in the cerebral cortex, as this is a region involved in behavioral processes disrupted in individuals with TBRS. This analysis confirmed broad reductions in mCA without profound global reductions in mCG (Figure 4C). To uncover site-specific changes in DNA methylation, we identified CG differentially methylated regions (DMRs) between sex-matched littermate pairs for both mutants. This analysis revealed thousands of hypo-methylated DMRs, with only a few hundred hyper-methylated DMRs in both mutants (Figure 4D-E). We detected 19,487 DMRs (196 hyper- and 19,291 hypo-DMRs) in the R878H mutant, while only 4,021 DMRs (215 hyper- and 3,906 hypo-DMRs) were observed in the P900L mutant, further indicating more dramatic methylation changes in the R878H mutant. Notably, however, hypo-DMRs called in one mutant were still generally also hypo-methylated in the other, demonstrating a broad concordance of effects with differing degrees of impact (Figure 4E-F). Interestingly, hyper-DMRs were not consistent between mutants suggesting these effects may be stochastic or secondary to DNMT3A disruption (Figure 4F). To further understand how these DMRs may be affecting transcription, we assessed their location in the genome and found that DMRs tend to fall in gene regulatory regions more than expected by chance, especially at CpG shores, enhancers, and regions that gain methylation during postnatal neuronal maturation (Adult DMRs) (Figure 4G). Thus, both mutants have numerous hypo-DMRs in critical gene regulatory regions, with R878H mutants displaying more severe effects than P900L mutants.

While DMR calling identifies a population of significantly altered mCG sites, DNMT3A mutants also display widespread changes in mCA that impact genomic regulatory regions but cannot be detected by DMR calling due to limitations in statistical power. Therefore, we next quantified overall levels of mCA and mCG across a number of regions of interest such as gene bodies, promoters, and enhancers. We found significant reductions in mCA and mCG at gene bodies and enhancers, and significant changes in mCA at promoters (Figure 4H-I). Regions that gain CG methylation during postnatal neuronal development (Adult DMRs) are particularly susceptible to DNMT3A disruption and have more dramatic changes (Figure 4I). The R878H mutant displayed significantly more reduction of mCA than the P900L across all regions, with more dramatic reductions of mCG compared to the P900L mutation at all sites examined (Figure 4H-I), thus highlighting the increased severity of the R878H mutation. These results indicate that both mutants exhibit loss of DNA methylation at critical genome regulatory regions that have the potential to affect enhancer activity and gene expression. Furthermore, these results demonstrate that the R878H mutation causes a more dramatic loss in neuronal methylation than the P900L mutation, which may drive an increase in phenotype severity.

### Altered enhancer activity corresponds with DNA methylation loss in P900L and R878H mutants

Enhancers are cis-regulatory elements important for controlling gene expression, and DNA methylation at both CA and CG sites control enhancer activity through effects on transcription factor binding and recruitment of methyl-DNA binding factors (Clemens & Gabel, 2020; Giacoman-Lozano et al., 2022; Kozlenkov et al., 2014). To determine if DNMT3A mutants have varied disruption of enhancers corresponding to their differential changes in DNA methylation, we examined enhancer activity by ChIP-seq of Histone H3 lysine 27 acetylation (H3K27ac), a histone modification correlated with active enhancers. We measured H3K27ac in the cortex of 8-week P900L and R878H mutants as well as WT littermates and used EdgeR to define changes at enhancers, allowing us to assess whether enhancers containing hypo- or hyper-CG DMRs were dysregulated. Enhancers containing a hypo-CG DMR showed significant increases in H3K27ac in the R878H mutant, and a trend towards upregulation in the P900L mutant (Figure 5A). To examine the relationship between mCA and enhancer activity more closely, we measured changes in mCA at the most significant 1% of upregulated and downregulated enhancers, allowing for comparison of both mutants using a similarly sized group of enhancers. This analysis revealed a more dramatic loss of mCA in upregulated enhancers compared to unchanged enhancers in both mutants (Figure 5B). These findings suggest that changes in mCA contribute to altered enhancer activity in both mutants, indicating a shared mechanism that may disrupt gene expression.

**Figure 5:**
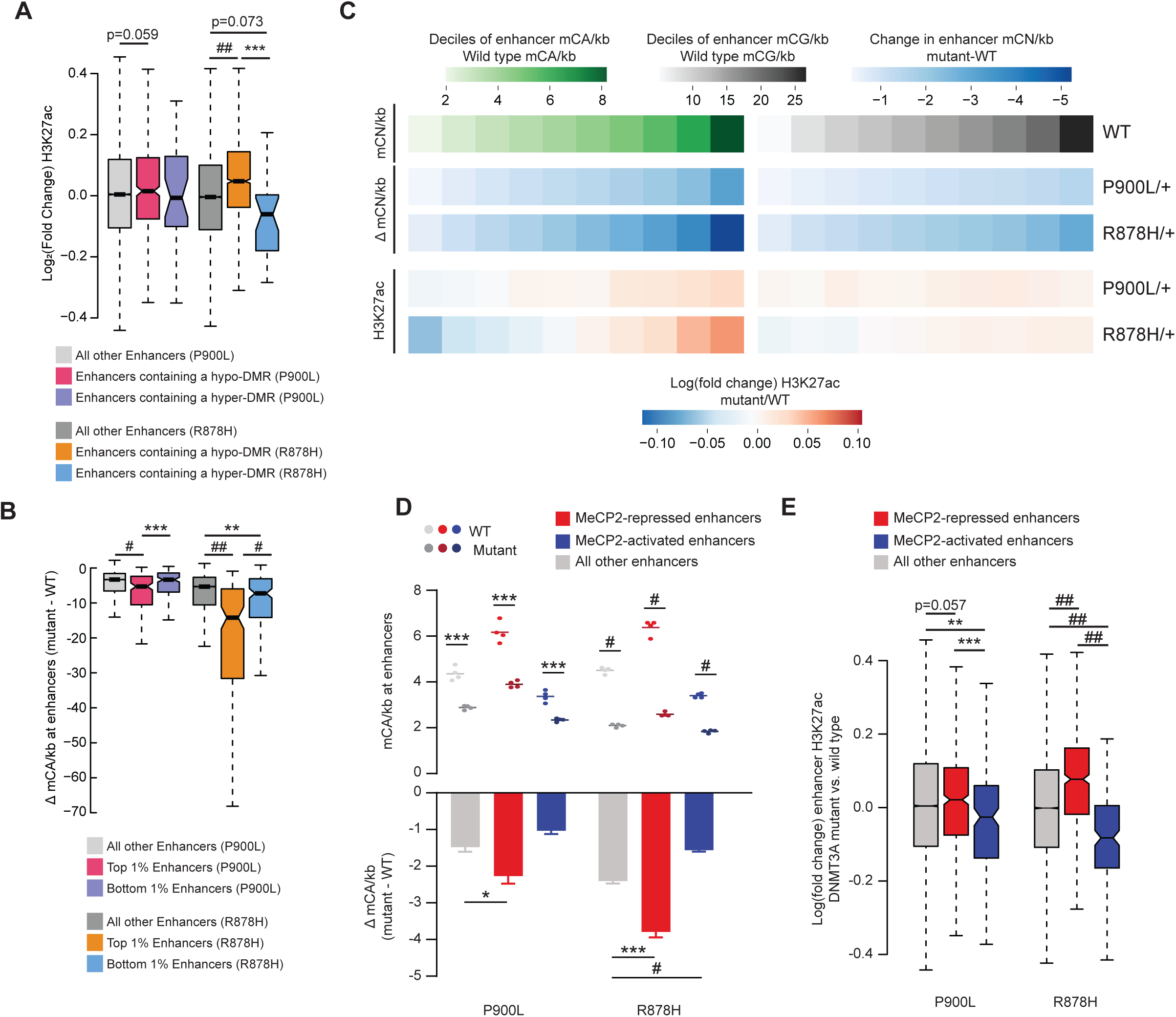
Methylation changes in DNMT3A mutants disrupt enhancer activity. (A) Log_2_ fold changes of H3K27ac at enhancers containing DMRs called in that mutant. (B) Change in mCA/kb (mutant – WT) for the top and bottom 1% of enhancers. The most significantly upregulated and downregulated enhancers were called between WT and mutants, and the average methylation loss at those enhancers was measured. (C) Genome-wide deciles of WT mCA or mCG sites per kilobase at enhancers, and the loss of methylation and fold change in H3K27ac at these sites in both P900L and R878H mutants. (D) Mean mCA sites per kilobase in DNMT3A mutants and their WT littermates (top), and the change in mCA sites per kilobase between mutants and WT littermates (bottom) at enhancers significantly dysregulated in MeCP2 mutants (Clemens *et al*., 2019). (E) Log_2_ fold changes in H3K27ac between mutants and WT littermates at enhancers significantly dysregulated in MeCP2 mutants (Clemens *et al*., 2019). Bar graphs indicate mean ± SEM. Notched box and whisker plots indicate median, interquartile, and confidence interval of median. All groups n=4, 2 male, 2 female. Detailed statistics, and sample sizes in Supplemental Table 1. * *p*<0.05; ** *p*<0.01; *** *p*<0.001; # *p*<0.0001; ## *p*<2 e^-10^

Our analysis of enhancers with significantly altered activity suggest more dramatic changes in the R878H mutant, however both mutants exhibited significant disruption of DNA-methylation broadly across the genome. Therefore, we next asked if there were broad changes in enhancer activity that correlate with genome-wide differences in DNA methylation and assessed if these changes differentially occur in P900L and R878H mutants. This analysis revealed that enhancers with a high density of WT mCA sites (mCA/kb) exhibited the largest corresponding loss of mCA in both mutants, and the greatest increases in H3K27ac (Figure 5C). Similarly, enhancers with low WT levels of mCA showed the smallest reductions in mCA and a decrease in relative H3K27ac. While this effect was observed in P900L mutants, the R878H mutation caused more dramatic changes in mCA corresponding with increased enhancer disruption. Genome-wide mCG-driven changes in enhancer activity are more subtle, implying that perhaps only a subset of enhancers with robust mCG differences are affected in either mutant (Figure 5C). This indicates that enhancers across the genome are sensitive to changes in mCA, and that DNMT3A-driven methylation changes at enhancers have the potential to contribute to gene expression changes.

DNMT3A and MeCP2 cooperate to regulate enhancer activity (Boxer et al., 2020; Clemens et al., 2019), therefore we next asked if enhancers most sensitive to MeCP2 disruption are differentially affected in the R878H mutant compared to the P900L mutant. Enhancers repressed by MeCP2 displayed greater loss of mCA in both mutants compared to other enhancers (Figure 5D). This increased loss of methylation is accompanied by a corresponding increase in H3K27ac for both mutants, with a more pronounced effect in the R878H mutant compared to the P900L mutant (Figure 5E). Thus, highly methylated enhancers are regulated by MeCP2, and loss of DNMT3A-dependent methylation at these enhancers causes overlapping disruption with loss of MeCP2, indicating shared effects between these two epigenetic regulators.

### Core disruption of growth genes across mutants with differential effects on synaptic and protein processing genes

DNA methylation and enhancer activity are critical for neurons to tune gene expression programs that contribute to development and function of the nervous system, and changes in gene expression can cause cellular and circuit disruption to drive disease phenotypes. We therefore used RNA-seq of cerebral cortex from 8-week mutant and WT littermate pairs to define alterations in transcriptional programs. P900L mutants displayed fewer significantly changed genes (Figure 6A: 892 up, 581 down) compared to R878H mutants (Figure 6B: 1,396 up, 1,326 down), mirroring the more severe epigenomic and behavioral effects in the R878H mutant. Gene expression changes in both mutants are concordant with a model of conditional deletion of DNMT3A from postmitotic neurons (Clemens et al., 2019), indicating that a number of these gene expression changes are related to the postnatal neuronal function of DNMT3A (Supplemental Figure 3A). Next, we used PANTHER to identify the enriched gene ontology (GO) terms in upregulated and downregulated genes to uncover the biological processes that may be most affected in these mutants. The genes upregulated in the P900L were enriched for critical neuronal functions related to synaptic transmission and axon guidance, such as cell-cell adhesion, and modulation and regulation of synaptic signaling, whereas there were no significantly enriched terms associated with the P900L-downregulated genes (Figure 6C). In contrast, the genes upregulated in the R878H were associated with protein folding, and the downregulated genes were associated with synaptic signaling, phospholipid translocation, and cell-cell recognition and assembly (Figure 6C). The distinct processes disrupted in these mutants may help explain the variable presentation of phenotypes; the P900L has more subtle changes in specific behavioral tasks perhaps driven by changes in synaptic and axonal genes, whereas the R878H mutant shows more widespread behavioral disruption corresponding with dramatic transcriptional changes involving fundamental biological processes such as protein folding and phospholipid translocation.

**Figure 6:**
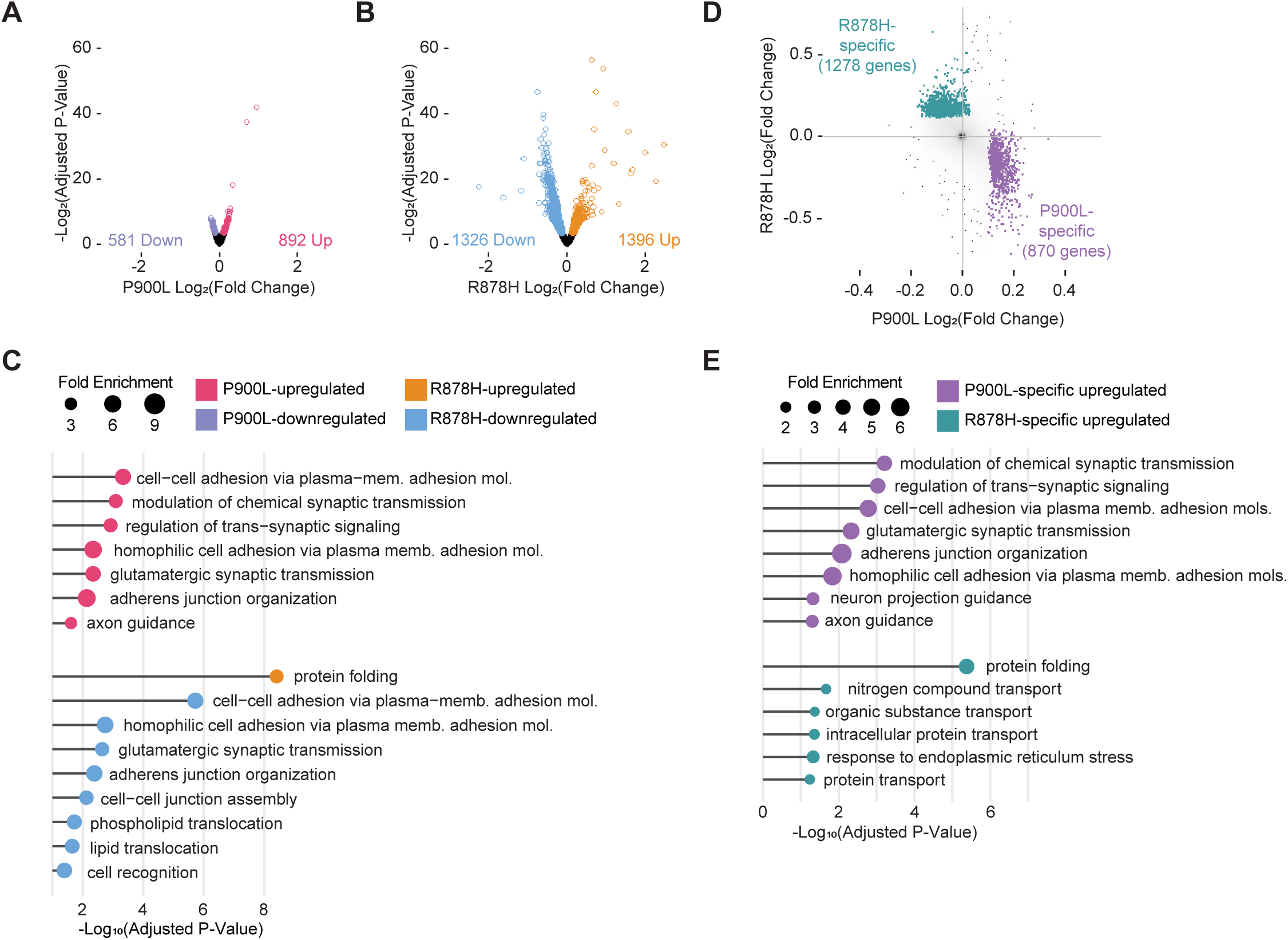
Mutation-specific changes in transcription indicate unique disruption in synaptic and protein processing gene networks. (A) Volcano plot of DESeq2 log_2_ fold changes in P900L vs. WT cortex. Genes reaching a significance of *p_adj_*<0.1 are indicated in purple and pink. (B) Volcano plot of DESeq2 log_2_ fold changes in R878H vs. WT cortex. Genes reaching a significance of *p_adj_*<0.1 are indicated in blue and orange. (C) Most significant PANTHER gene ontology (biological process) terms enriched in each differentially expressed gene list. No significant terms were identified in the P900L- downregulated gene list. (D) P900L- and R878H-specific upregulated gene sets indicated in purple and teal. Specific genes are defined as those that are significantly upregulated in one mutant, and either significantly unchanged (nominal p-value > 0.5) or downregulated in the other (fold change < 0). (E) Most significant PANTHER gene ontology (biological process) terms enriched in P900L- specific and R878H-specific upregulated gene lists.

To more directly assess the transcriptomic differences that could lead to distinct phenotypes between mutants, we defined mutant-specific genes by identifying genes upregulated in one mutant and either unchanged or downregulated in the other mutant (Figure 6D), and again used PANTHER to identify enriched gene ontology terms. We focused first on upregulated gene lists, as these may be the most direct targets from loss of DNA methylation. The P900L-specific upregulated genes again were primarily enriched for synaptic and axonal projection processes, whereas the R878H-specific upregulated genes were enriched for protein folding and transport terms (Figure 6E). The R878H-specific downregulated terms were primarily related to glutamatergic synaptic transmission and cell-cell adhesion and organization, whereas P900L-specific downregulated genes were not associated with any GO terms (Supplemental Figure 3B-C). This further suggests that transcriptional disruption in the P900L affects fine-tuned and sensitive neuronal processes, whereas dysregulated genes in the R878H are potential indicators of more dramatic and widespread cellular distress.

While leveraging the transcriptional and phenotypic differences between mutants offers insight into which gene sets contribute to distinct phenotypes, identifying the shared effects across multiple DNMT3A mutant models can identify central biology driving common disease phenotypes. Therefore, we leveraged an existing disease-relevant cortical dataset from the DNMT3A deletion mouse model and analyzed it together with our new transcriptomic data to identify the shared disruption in DNMT3A mutant models. We used our data from the P900L/+ (n=7/genotype; 4 male, 3 female) and R878H/+ (n=7/genotype; 4 male, 3 female) mutations, and gene expression data from a heterozygous KO mouse (n=7/genotype; 4 male, 3 female) (Christian et al., 2020), to create an aggregate dataset of heterozygous germline DNMT3A models. We then used DESeq2 to compare WT and mutant gene expression between sex-matched littermate pairs (design = ∼ pair + group; contrast by group to identify WT vs. mutant effects) thus identifying high-confidence TBRS-associated differentially expressed genes (Figure 7A). This analysis identified 228 upregulated and 160 downregulated genes that show concordant up- and down-regulation across mutant strains (Figure 7B). TBRS-upregulated genes are enriched for processes such as cellular and developmental growth, axon extension, and neural crest cell migration and no terms were significantly enriched in the downregulated genes (Figure 7C). The dysregulated genes in these pathways may be critical in driving the overgrowth and behavioral phenotypes identified in individuals with DNMT3A disorders and represent strong candidates for future cellular and therapeutic studies.

**Figure 7:**
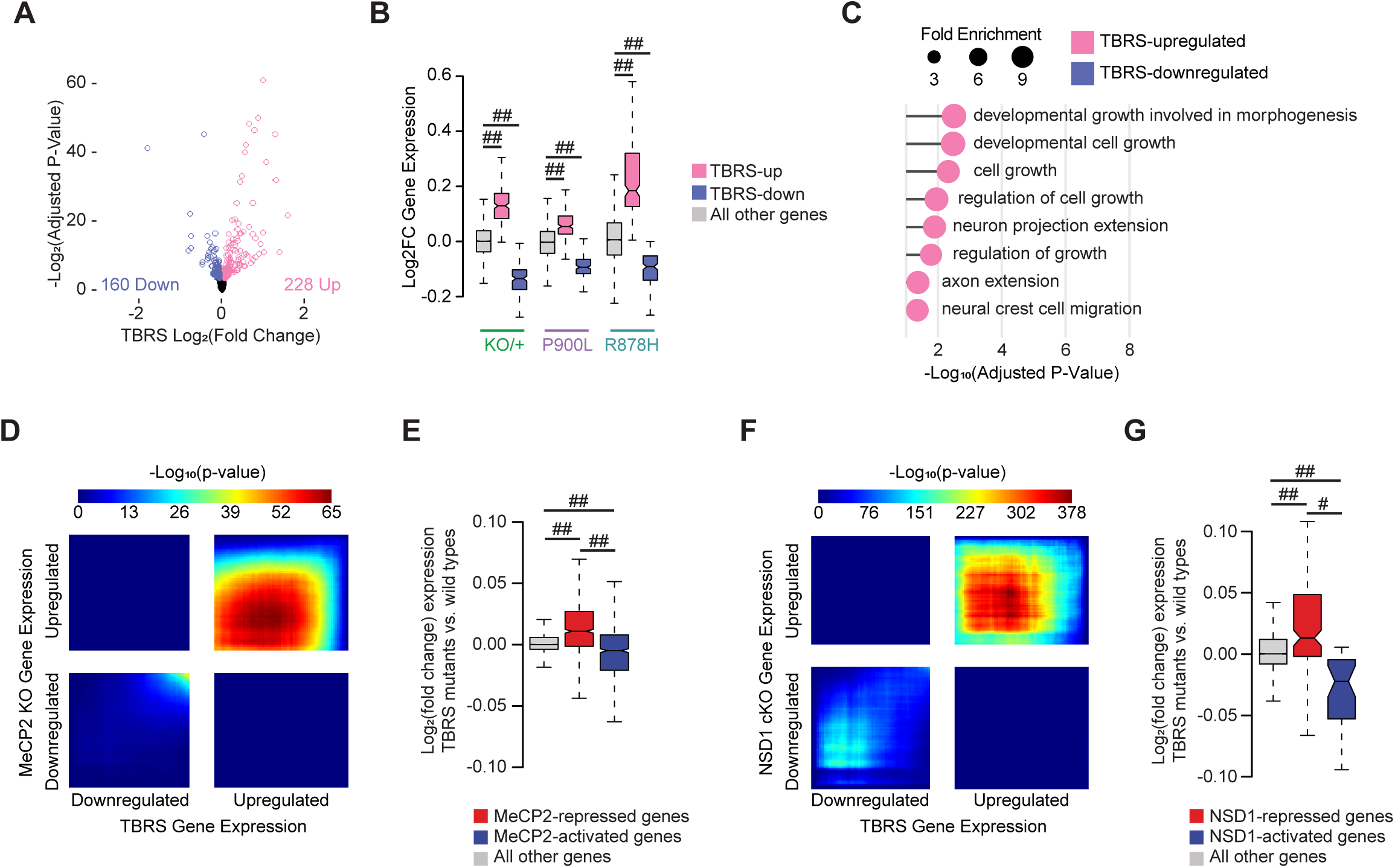
Shared transcriptional changes across DNMT3A mutants indicate disruption of growth and synaptic processes. (A) Volcano plot of DESeq2 log_2_ fold changes from DNMT3A mutant vs. WT littermate analysis between littermate paired data from DNMT3A^KO/+^ (Christian *et al*., 2020), DNMT3A^R878H/+^, and DNMT3A^P900L/+^ datasets (design = ∼ pair + group; contrast by group). Genes reaching a significance of *p_adj_*<0.1 are indicated in blue and pink (B) Log_2_ fold changes of gene expression within each mutant (KO/+, P900L, and R878H) of genes defined as differentially expressed in the combined TBRS-mutant analysis. (C) Most significant PANTHER gene ontology (biological process) terms enriched in the TBRS differentially expressed gene lists. No significant terms were identified in the TBRS-downregulated gene list. (D) Rank-rank hypergeometric overlap (RRHO) of transcriptome-wide gene expression changes in the cerebral cortex of TBRS mutants versus MeCP2 KO mice (Clemens *et al*., 2019). (E) Log_2_ fold changes in the TBRS mutants at genes significantly disrupted in MeCP2 mutants (Clemens *et al*., 2019). (F) RRHO of transcriptome-wide gene expression changes in the cerebral cortex of TBRS mutants versus NSD1 conditional KO mice (Hamagami *et al*., 2023). (G) Log_2_ fold changes in the TBRS mutants at genes significantly disrupted in NSD1 cKO cortices (Hamagami *et al*., 2023). Notched box and whisker plots indicate median, interquartile, and confidence interval of median with significance from Wilcox Rank Sum test shown. Detailed statistics, and sample sizes in Supplemental Table 1. # *p*<0.0001; ## *p*<2 e^-10^

### Genes disrupted in TBRS models are shared across disorders that impact the neuronal methylome

Multiple neurodevelopmental diseases are caused by mutations in genes associated with the neuronal methylome (e.g., DNMT3A (Tatton-Brown et al., 2018), MeCP2 (Tillotson et al., 2021), NSD1 (Hamagami et al., 2023)), so we next asked if transcriptomic disruption is shared between multiple models of neurodevelopmental disorders. Previous work has shown that deletion of MeCP2 and DNMT3A have overlapping gene expression patterns (Christian et al., 2020; Clemens et al., 2019; Lavery et al., 2020), and we have established that MeCP2- regulated enhancers are similarly disrupted in the P900L and R878H mutants, therefore we next asked if the core TBRS-dysregulated genes overlap with the gene expression changes observed in the MeCP2 KO. We performed a Rank-Rank Hypergeometric Overlap (RRHO) analysis (Cahill et al., 2018) to measure transcriptome-wide correspondence between the TBRS-mutant models and the MeCP2 KO and found significant overlaps in the concordant quadrants (Figure 7D). Additionally, genes significantly dysregulated in MeCP2 mutants are correspondingly disrupted in the TBRS mutants (Figure 7E, Supplemental Figure 3D). Transcriptional overlap between this consensus TBRS dataset and the MeCP2 KO further supports a shared molecular etiology between mutation of DNMT3A, which methylates the neuronal genome, and MeCP2, which binds that methylation to repress transcription of genes.

Overlapping clinical phenotypes or shared biological pathways can be used to suggest other important candidate regulators of DNMT3A and the neuronal methylome. One such candidate is NSD1, a histone methyltransferase associated with Sotos Syndrome (Saugier-Veber et al., 2007; Tatton-Brown et al., 2005). A significant number of patients with overgrowth and intellectual disability phenotypically similar to TBRS patients have mutations in NSD1 (Tatton-Brown et al., 2017), and emerging studies have also demonstrated that NSD1 deposits H3K36me2 to direct DNMT3A to establish methylation at key genomic regions in neurons (Hamagami et al., 2023). This led us to ask if there are shared effects between NSD1 mutants and the core gene dysregulation we identified in TBRS models. RRHO comparison of cortical genes dysregulated in an NSD1 conditional KO model and aggregate TBRS effects indicates a highly significant transcriptome-wide concordance (Figure 7F), and genes identified as significantly dysregulated in the NSD1 mutant are similarly dysregulated in the TBRS mutants (Figure 7G, Supplemental Figure 3E). Interestingly, for both MeCP2 and NSD1 comparisons, overlaps were more pronounced in shared upregulated genes, suggesting these are the direct effects of the pathway. Downregulated genes across models were more unique, suggesting they may be more stochastic and less central to the shared phenotypes of the three syndromes. Together these results indicate that the core sets of genes driving DNMT3A-disorders are shared with models of Rett Syndrome and Sotos Syndrome. This reflects a biological convergence across multiple disorders, indicating the neuronal mCA-pathway regulating gene expression may be a useful target in future studies of potential therapeutics for all three disorders.

## Discussion

Neurodevelopmental disorders (NDDs) often present with varied phenotypes and numerous comorbidities, and the molecular mechanisms driving this spectrum of phenotypic heterogeneity have not been clearly identified. Additionally, a substantial number of mutations identified in some NDD-associated genes are missense rather than stop-gains (e.g., *KIF1A, MEFC2, CHD3, PTEN, GRIN2B, DNMT3A*), and the effects of these diverse mutations are not fully understood (Wang et al., 2020). Here, we studied missense mutations in DNMT3A to investigate the origins of clinically diverse phenotypes within one causative locus, ranging from ASD to severe intellectual disability. We identified behaviors that indicate varied severity of alleles and linked these changes to differential disruption of neuronal methylation and transcription. Through this work, we not only identified the core set of phenotypes and shared genes that are central to DNMT3A-disorders, but also defined allele-specific gene networks and cellular processes that may underlie the spectrum of phenotypes. Furthermore, we detected transcriptional overlap between core DNMT3A gene expression effects and disruption of MeCP2 and NSD1, highlighting a potential point of convergence in disease etiology and therapeutic intervention. In this study, we generated a clinically-relevant germline mutation that represents the larger class of “typical” missense mutations in the methyltransferase domain of DNMT3A. Through our analysis, we identified skeletal development and obesity phenotypes that are consistent across multiple DNMT3A mutations. The increase in long-bone length shared between the P900L mutation and other mutants underscores the importance of DNMT3A in skeletal development and growth. P900L mutants also exhibit similar increases in body fat compared to other DNMT3A mutants (Christian et al., 2020; Smith et al., 2021; Tovy, Reyes, et al., 2022), and we expand these observations by demonstrating that progressive increase in fat mass can occur without changes in feeding behavior or substantial decreases in exploratory behaviors. This suggests other metabolic or cellular processes may be responsible for obesity in DNMT3A mutants, and further highlights the potential mechanism proposed by Tovy et al. that DNMT3A mutations may cause expansion of adipocyte progenitors (Tovy, Reyes, et al., 2022). These findings reinforce the importance of DNMT3A in skeletal development and provide important context supporting the role of DNMT3A in obesity.

Our analysis of skull size and shape demonstrates that the P900L mutation does not exhibit changes in skull morphology, which is similar to other mouse models of TBRS (Christian et al., 2020; Smith et al., 2021) but does not phenocopy the human disorder. We also identified reductions in brain volume shared across multiple mutants. The lack of brain overgrowth in mice could contribute to the lack of differences in skull size and shape, as skull development is sensitive to changes in brain volume (Bartholomeusz et al., 2002). Evolutionary differences in growth regulation between humans and mice may underlie these phenotypic distinctions, suggesting that other approaches, such as using human cellular models, may be necessary to identify the mechanisms leading to human brain overgrowth. Notably, the reduction in brain size is shared between multiple mouse mutations, suggesting that the cellular processes leading to decreased brain volume (e.g., changes in cell counts, cell size or dendritic arborization) are highly sensitive to DNMT3A disruption. Furthermore, this may represent phenotypic overlap between DNMT3A- and MeCP2-mutants that reflects the similar transcriptional disruption that we observe between these models. This is further supported by the lack of brain volume differences at P10, a timepoint prior to widespread MeCP2 expression in the brain.

Humans with DNMT3A mutations range in clinical diagnoses from ASD to severe intellectual disability, and our characterization of the P900L model allowed us to identify behavioral domains with similar phenotypic heterogeneity. Previous work demonstrated that heterozygous loss of DNMT3A in mice causes reduced exploration and increased anxiety-like behaviors (Christian et al., 2020), and R878H mutant mice have more dramatic reductions to exploratory behavior and disruption of motor coordination (Smith et al., 2021). In contrast, the P900L mutant has no motor, exploratory, or anxiety-like changes, indicating that these phenotypes are not ubiquitous across all mouse models of DNMT3A disruption, and instead may be representative of the phenotypic heterogeneity of DNMT3A disorders. We also identified severe alteration of social hierarchies in the R878H mutant that are not observed in the P900L mutant. These findings clearly demonstrate differences in phenotype severity and define differential phenotypes that we can compare to altered epigenomic and transcriptional effects in these models.

We expanded DNMT3A-phenotypes in mice by assessing behaviors associated with ASD and identified disruption of communication and tactile discrimination shared across multiple models. Altered tactile discrimination is an emerging phenotype across multiple ASD models (Orefice et al., 2019; Orefice et al., 2016), indicating a potential mechanism contributing to behavioral disruption, and highlighting the importance of DNMT3A in sensory processing. Our study also confirms that neonatal ultrasonic vocalizations are reproducibly sensitive to DNMT3A disruption, as demonstrated by the reduction in number of calls in the P900L mutants, R878H mutants, and in a heterozygous KO model (Christian et al., 2020). This work establishes that communication and tactile discrimination deficits are consistent and robust across DNMT3A models, suggesting that these measures are a promising focus for future work testing therapeutics or identifying cellular mechanisms contributing to disruption.

Behavioral differences in individuals with DNMT3A mutations and the numerous functions of DNMT3A in nervous system development indicate importance of this protein in the brain. Our work here defines how disease-associated missense mutations affect the neuronal epigenome *in vivo*, thus beginning to uncover mechanisms driving behavioral phenotypes and nervous system disruption. We found that the P900L mutation causes a 50% reduction of mCA which mimics a heterozygous KO, supporting the hypothesis that mCA levels are a sensitive readout of DNMT3A function. In contrast, the R878H mutation causes greater than 50% loss of mCA, demonstrating that it drives more dramatic effects than other mutations. While our results do not shed light on the exact mechanism leading to this effect, this *in vivo* result supports studies in the blood lineage indicating that R878H mutation is dominant-negative (Russler-Germain et al., 2014). In addition to these insights, our study illustrates the sensitivity of global mCA levels to different DNMT3A mutations and suggests that even weak loss of function mutants may have alterations in global mCA levels. Differences in allele-severity are further reflected by the increase in number of DMRs in the R878H mutant compared to the P900L mutant, and we demonstrate that these methylation differences overlap with key genomic regulatory elements such as gene bodies and enhancers. Notably, we found increased enhancer disruption in the R878H mutant that corresponds to larger changes in mCA at enhancers. This enhancer effect is similar to observations in MeCP2 mutants, and we demonstrate that DNMT3A mutants have concordant disruption of enhancers regulated by MeCP2. Together, these results are the first to demonstrate how disease-associated missense mutations in DNMT3A differentially disrupt numerous neuronal epigenomic processes and suggest a molecular mechanism driving the spectrum of phenotypic severity.

Our work defined mutation-specific gene expression changes to gain insights into the cellular disruptions and biological pathways that may be driving the spectrum of disease phenotypes. The P900L mutation causes disruption of fine-tuned neuronal genes related to synaptic function and axonal guidance, suggesting that synapses, axon projections, and circuit connectivity may be disrupted in mutants. It is possible that these transcriptional changes are contributing to the reduction in FA measured in the P900L corpus callosum, indicating potential disruption of long-range axonal projections in this model. The R878H mutation caused more extensive transcriptomic disruption, altering gene networks involved in key cellular processes such as protein folding and molecular transport. These allele-specific transcriptomic effects identify cellular mechanisms that may underlie the unique behavioral phenotypes and provide compelling candidates for future work on distinct cellular- and circuit-level effects in DNMT3A disorders.

Our characterization of transcriptional disruption in multiple models allowed us to define the core sets of neuronal genes most sensitive to DNMT3A mutation that may contribute to TBRS pathology. Genes consistently dysregulated across DNMT3A mutants suggest cellular mechanisms that may be disrupted in TBRS. Upregulation of NDD-associated genes such as the Semaphorin family (*Sema3b, Sema3e, Sema4a, and Sema5a*) and *Tbr1* suggest changes in axon guidance and migration may contribute to TBRS pathology, and these effects could be involved in disruption of ultrasonic vocalizations in mice (Co et al., 2022; Duan et al., 2014; Fazel Darbandi et al., 2018, 2020; T. N. Huang et al., 2014; Sollis et al., 2022; Zhao et al., 2018). Changes in sensory neurons contributing to NORT phenotypes in DNMT3A mutants may be related to disruption of *Etv4*, *Myocilin*, and *Begain*, as these genes are important for the proper development, growth, and myelination of peripheral neurons (Katano et al., 2016; Kwon et al., 2013; Ríos et al., 2022; Smit et al., 2005; Yao et al., 1996). Finally, downregulation of *Sox21* and *Gabrg1* in DNMT3A mutants suggest potential changes in GABAergic interneurons and precursors (Makrides et al., 2018; Polan et al., 2014) which may be playing a role in developmental delay (Williams et al., 2022). Importantly, several classes of interneurons have high global levels of mCA (Mo et al., 2015), offering a potential mechanism to explain why interneurons may be uniquely vulnerable to loss of DNMT3A function. Together, this work defines the gene sets and processes that are most susceptible to DNMT3A disruption and provides insights into potential biological processes and cell types that could contribute to disease.

Finally, our study has detected transcriptional convergence between core gene dysregulation in TBRS models and mutations in other proteins in the neuronal-methylome pathway, supporting potential functional links between Sotos syndrome, TBRS, and Rett Syndrome. The transcriptional similarities between disruption of DNMT3A and other epigenetic regulators highlights the importance of this pathway for neuronal gene regulation and indicates a therapeutic point of convergence across an entire class of NDDs.

## STAR METHODS

### Key Resources Table

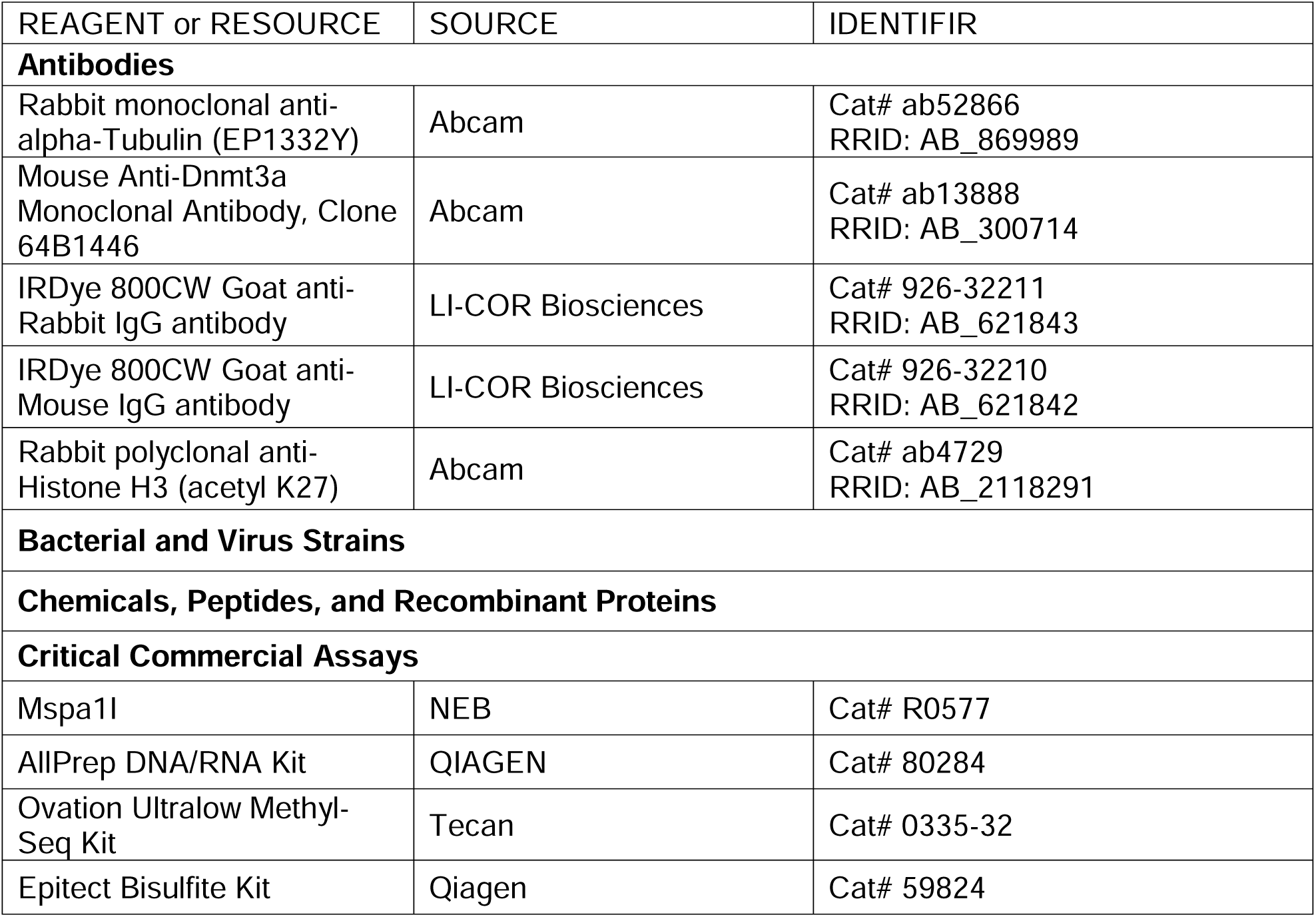
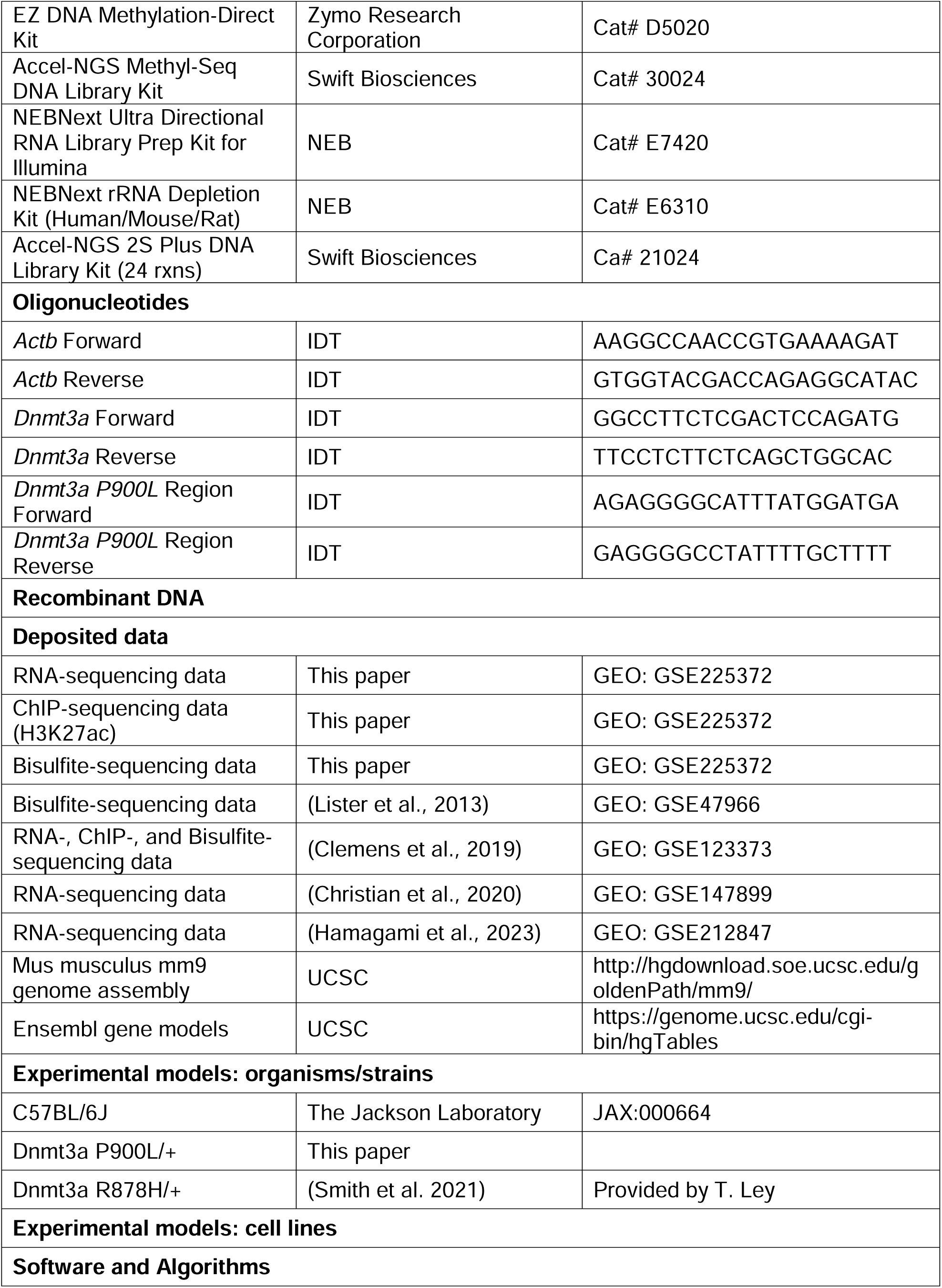
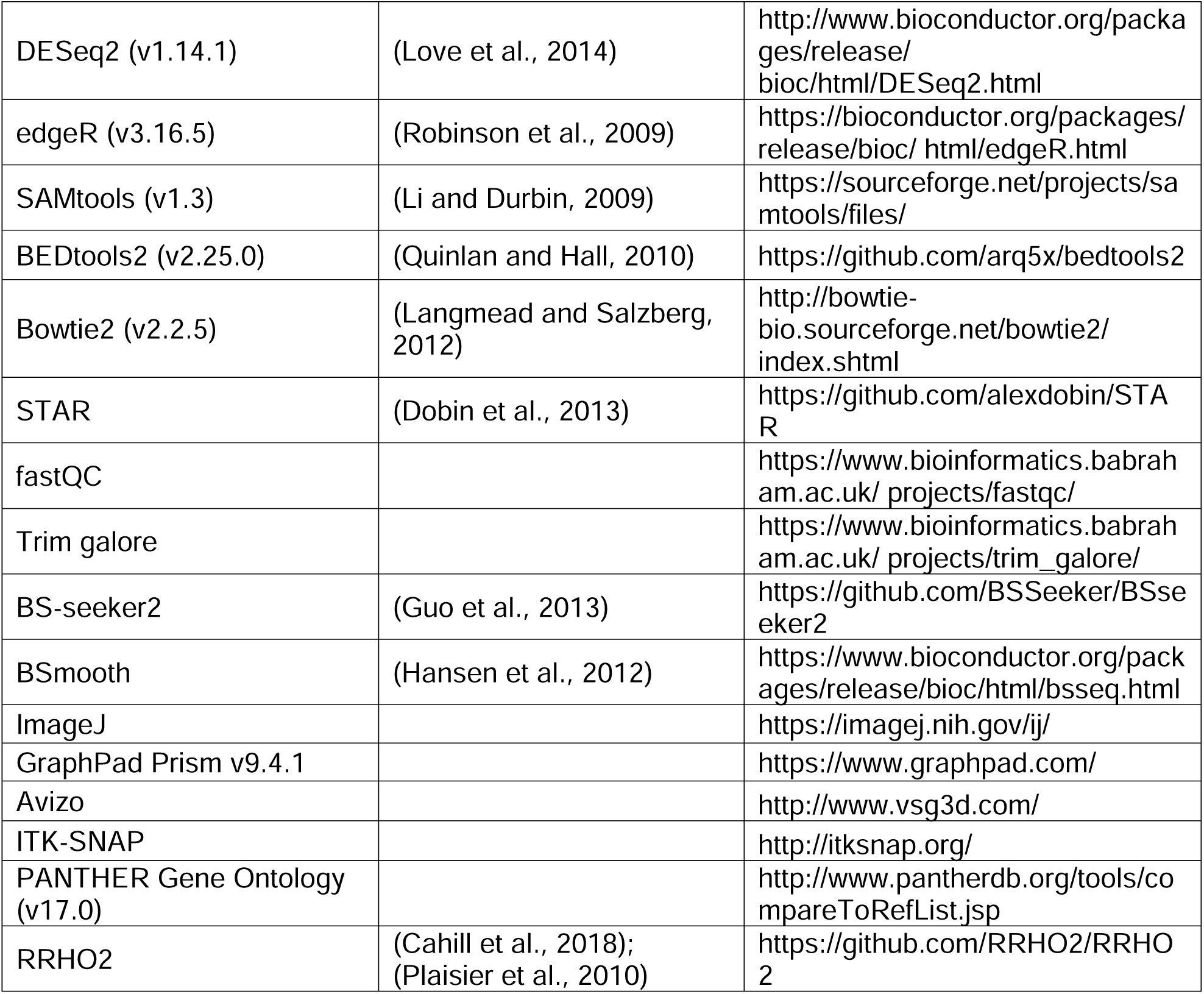

### Contact for Reagent and Resource Sharing

Requests for reagents and resources should be directed toward the Lead Contact, Harrison Gabel.

### Experimental Model and Subject Details

#### Animal Husbandry

All animal protocols were approved by the Institutional Animal Care and Use Committee and the Animal Studies Committee of Washington University in St. Louis, and in accordance with guidelines from the National Institutes of Health (NIH). Mice were housed in a room on a 12:12 hour light/dark cycle, with controlled room temperature (20-22°C) and relative humidity (50%). Home cages (36.2 x 17.1 x 13 cm) were individually ventilated and supplied with corncob bedding and standard laboratory chow (PicoLab Irradiated Rodent Diet 5053) and water unless otherwise specified. For experiments of progressive weight gain, male and female animals (P900L n=18, 8 male, 10 female; WT n=24, 12 male, 12 female) were given free access to the Tekkad High Fat Diet (Envigo; TD.88137; 42% Calories from Fat) instead of standard laboratory chow from 10-30 weeks of age. During this time, mice were weighed weekly. At 30 weeks of age, mice were single housed, and food was weighed every two days for a total of six days (3 timepoints) to measure food consumption. Unless otherwise specified, all mice were group-housed and adequate measures were taken to minimize animal pain or discomfort.

#### Transgenic animals

The DNMT3A P900L mouse model was generated using single guide RNAs (sgRNAs) to create a C→T substitution at chr12:3,907,719 (GRCm38/mm10 assembly). This mutation changed the proline CCG codon into a leucine CTG codon (Supplemental Figure 1A). sgRNAs were cloned into the pX330 Cas9 plasmid (Addgene), and then transfected into N2A cells. Validation was done using the T7 enzyme assay by the Washington University School of Medicine Transgenic Vectors Core. sgRNAs were transcribed *in vitro* using MEGAShortScript (Ambion), and Cas9 mRNA was *in vitro* transcribed, G-capped and poly-A tailed using the mMessageMachine kit (Ambion). mRNA of the sgRNA and Cas9 were then injected into hybrid C57Bl/6J x CBA fertilized eggs at the mouse genetics core at Washington University School of Medicine. Founders were deep sequenced at expected cut sites to identify which alleles were present, and deep sequencing analyses of four kilobases surrounding the targeted region was used to ensure no off-target recombination events occurred. Founders were then crossed to C57BL6/J females (JAX Stock No. 000664) for 5-10 generations before experimental analysis.

To generate experimental animals, Dnmt3a^R878H/+^ (R878H) or Dnmt3a^P900L/+^ (P900L) male mice were crossed with C57BL6/J females (JAX Stock No. 000664). R878H and P900L females were not used for breeding to avoid social differences in mothering from mutant dams. Mice were genotyped with ear-, tail-, or toe- DNA by PCR for either R878H or P900L mutations. Mice were weighed at a variety of timepoints to assess growth.

#### Method Details P900L Genotyping

To genotype for the P900L mutation, ear-, tail-, or toe- DNA was amplified using primers designed around the P900L mutation (F:AGAGGGGCATTTATGGATGA, R: GAGGGGCCTATTTTGCTTTT). The 706bp PCR product could then be Sanger Sequenced (Supplemental Figure 1A) or digested using Mspa1I for an extended 3-hour digestion time followed by the standard heat-shock inactivation. The wild-type sequence is susceptible to restriction enzyme digestion, leaving a 285bp and 421bp fragments, whereas the P900L mutation is not digested and will remain at 706bp (Supplemental Figure 1B).

#### Bone length measurements

We chose to quantify long bones that may directly relate to the height phenotype seen in patients. Femurs were dissected from 30–35-week-old mice (P900L n=18, 10 male, 8 female; WT n=16, 9 male, 7 female) and scanned using a Faxitron Model UltraFocus100 Digital Radiography system at the Washington University Musculoskeletal Research Center. Image analysis was done using Faxitron Vision Software (Version 2.3.1). When analyzed with a 2-way ANOVA, there was no significant sex effect. Bone lengths were also measured from dissected femurs using a Vernier caliper, which yielded similar results (data not shown).

#### Craniofacial morphological analyses

A total of 16 sex-matched littermate paired mice (P900L n=8, 4 male, 4 female; WT n=8, 4 male, 4 female) at 30-35 weeks of age were fixed with intracardiac perfusions of 4% paraformaldehyde. Whole mouse heads were scanned using a Scanco µCT40 machine at the Musculoskeletal Research Center at Washington University in St. Louis. Image processing was performed as previously described (Christian et al., 2020; Hill et al., 2013). Briefly, CT images were converted to 8-bit and surface reconstructions were acquired in Avizo (http://www.vsg3d.com/). 35 landmarks were collected from surface reconstructions of the cranium and mandible using Avizo. Principal components were identified from generalized Procrustes analysis in Geomorph package in R and Morphologika software as previously described (Hill et al., 2013). To identify specifically altered linear distances, landmark coordinates were natural log-transformed and analyzed with linear regression using Euclidean Distance Matrix Analysis (EDMA).

#### EchoMRI to measure body composition

Fat and lean mass measures of live WT and P900L mice were measured with whole-body quantitative magnetic resonance using an EchoMRI Body Composition Analyzer at the Washington University Diabetes Research Center. Experiments were performed as previously described (Nixon et al., 2010). Briefly, animals of 30-35 weeks of age (P900L n=15, 8 male, 7 female; WT n=17, 10 male, 7 female) were placed in a plastic cylinder tube with a solid insert to limit movement. Signal in response to a low-intensity electromagnetic field was used to measure the relaxation of spin curves, allowing for the quantification of fat and lean tissue volume. Canola oil was used to standardize measurements between different recording days.

#### Magnetic Resonance Imaging (MRI) acquisition and Diffusion Tensor Imaging (DTI) analysis

A total of twenty-four animals were used for P900L experiments (WT n=12, 6 males, 6 females; P900L n=12, 6 males, 6 females), and twenty-four animals for R878H experiments (WT n=12, 6 males, 6 females; R878H n=12, 6 males, 6 females). Imaging and analysis were performed as described previously (Chen et al., 2021). In brief, isoflurane-anesthetized animals were scanned with a small-animal MR scanner built around an Oxford Instruments 4.7T horizontal-bore superconducting magnet equipped with an Agilent/Varian DirectDrive^TM^ console. Data were collected using a laboratory-built actively decoupled 7.5-cm ID volume coil (transmit)/1.5-cm OD surface coil (receive) RF coil pair. Mouse respiratory rate and body temperature (rectal probe) were measured with a Small Animal Instruments (SAI, Stony Brook, NY) monitoring and gating unit.

T2-weighted trans-axial images (T2W), collected with a 2D fast spin-echo multi-slice (FSEMS) sequence, were used for structural and volumetric analyses. Diffusion Tensor Imaging (DTI), which measures the directional movement of water along and perpendicular to axons (fractional anisotropy: FA), provided a measure of white-matter track integrity. DTI data were collected using a multi-echo, spin-echo diffusion-weighted sequence with 25-direction diffusion encoding, max b-value = 2200 s/mm^2^, as described previously (Chen et al., 2021). Two echoes were collected per scan, with an echo spacing of 23.4 ms, and combined offline to increase signal-to-noise ratio (SNR), resulting in a SNR improvement of ∼1.4x compared with a single echo.

DTI data were analyzed as described previously (Chen et al., 2021) according to the standard MR diffusion equation (Stejskal & Tanner, 1965) using purpose-written MatLab software. Eigenvalues (λ_1_, λ_2_, λ_3_) corresponding to the diffusion coefficients in three orthogonal directions, and parametric maps of apparent diffusion coefficient (ADC), axial diffusion (D_axial_), radial diffusion (D_radial_), and fractional anisotropy (FA) were calculated according to standard methods (Basser & Pierpaoli, 2011; Mori S & Tournier J-D, 2014). Parametric maps were converted into NIfTI (.nii) files for inspection and segmentation using ITK-SNAP (www.itksnap.org). Segmentation was performed blinded to strain, sex, or genotype, and consistency was assessed by re-segmenting blinded data files.

#### Behavioral Analyses

Mice for behavioral testing were housed in mixed genotype home cages with 2-5 animals per cage, and all tests were performed during the light cycle. All experimenters were female and were blinded to genotype during testing. For increased experimental rigor and reproducibility, we used separate cohorts of mice to ensure quality and consistency in any observed phenotypes. Adult testing was performed when mice were 2-4 months of age.

##### Maternal Isolation-Induced Ultrasonic Vocalizations

Pup ultrasonic vocalization (USV) measurements were performed to assess early social communicative behavior as previously described (Chen et al., 2021). Ninety-seven animals were used for P900L experiments (n=47 WT, 19 males and 28 females; n=50 P900L, 30 males and 20 females), and ninety-two animals were used for R878H experiments (n=51 WT, 24 males, 27 females; n=41 R878H, 19 males and 22 females). Recordings were done at postnatal days 5, 7, and 9. In brief, adults were removed from the nest and home-cages were placed in a warming box (∼33°C) 10 minutes before recording began. Body temperature was recorded immediately before placing pups in a dark, enclosed chamber for 3-minute recordings. Following the USV recording, pups were weighed and returned to their nest. Frequency sonograms were prepared and analyzed in MATLAB as previously described (Chen et al., 2021; Christian et al., 2020). Within-subjects repeated measures ANOVA were used to assess significance, and no significant differences occurred between sexes for any vocalization measures, therefore data were combined between sexes.

##### Marble burying

WT (n=13; 8 male, 5 female) and litter matched P900L (n=13; 8 male, 5 female) mice were used for marble burying as previously described (Christian et al., 2020). In brief, 8-week-old mice were placed in a transparent enclosure (28.5 cm x 17.5 cm x 12 cm) with clean aspen bedding and 20 dark blue marbles evenly spaced in a 4 x 5 grid on top of the bedding. Animals were allowed to explore freely for 30 minutes, and the number of buried marbles were counted every 5 minutes by two independent blinded observers. Marbles were considered “buried” if they were at least two-thirds covered by bedding. Enclosure and marbles were cleaned thoroughly between animals. Data was analyzed with a within-subjects repeated measured ANOVA, and no sex effect was observed so data was combined between sexes.

##### 3-Chamber social approach

Eighteen litter-matched animals that were 10-12 weeks old were used in the 3-chamber social approach assay (P900L n=9, 5 male, 4 female; WT n=9, 4 male, 5 female) as previously described (Manno et al., 2020). Briefly, mice were acclimated to a clear acrylic rectangular apparatus (60 cm x 40.5 cm), which was separated into three chambers by walls with sliding doors (6 cm x 6 cm). The apparatus was placed in an isolated, quiet room with low light (270 lux) to minimize stress. Both side chambers contained an inverted cup. Testing consisted of three 10-minute phases: during the first phase, the mouse freely explored all chambers, in the second phase a conspecific mouse was added to one of the cups (mouse vs. object), and in the third phase a novel conspecific was added to the remaining empty cup (novel vs. familiar). During all phases, the test mouse was allowed to freely explore, and all stimulus mice were sex-matched conspecifics. A digital video camera was used to record sessions, location of mice in the apparatus was analyzed. Between experimental animals, 70% ethanol was used to clean the apparatus. As mice rapidly habituate to this task (Manno et al., 2020), only the first 5 minutes of each phase was used for analysis.

##### Social Dominance Tube Test

Tube test was conducted to assess social hierarchy behavior as previously described (Chen et al., 2021). For P900L experiments, 94 animals were used (n=47 WT, 24 males and 23 females; n=47 P900L, 24 males and 23 females) across three experimental cohorts, and one cohort of 34 mice was used for R878H experiments (n=17 WT, 9 males, 8 females; n=17 WT, 9 males, 8 females). In brief, mice were allowed to learn to traverse the clear acrylic tube apparatus on days 1 and 2 of the task. On days 3-5, sex-matched pairs of WT and mutant mice were tested on dominance bouts, avoiding cage mate pairings. A new WT-mutant pairing was used each day, allowing for three distinct matchups for each animal. During bouts, animals were allowed to enter the tubes while separated from each other with an acrylic divider. A bout begins when the divider was removed and concluded when one mouse fully backed out of the tube or when 2 minutes passed. The animal remaining in the tube was considered the winner of the bout (dominant) and the animal that exited the tube was the loser (submissive). Active wins were defined as the winner pushing the other animal from the tube, whereas passive wins were defined as the winner refusing to move and the loser backing out of the tube. The tube was cleaned with a 0.02% chlorhexidine solution between bouts. Bout recordings were scored by a blinded observer. A two-tailed binomial test was performed on numbers of bouts won, with a null hypothesis that 50% of bouts would be won by each genotype.

##### Novel Object Recognition – Tactile

Novel Object Recognition-Tactile (NORT) was used to measure general and tactile associative memory adapted from previous work (Orefice et al., 2019; Orefice et al., 2016). Briefly, the task consisted of five consecutive days including two initial habituation trials, NORT testing, a third habitation trial, and NOR testing. During habituation trials, mice were allowed to freely explore the empty acrylic apparatus (26 x 26 cm or 40 x 40 cm) for 10 minutes under white light (75-100 lux). During NORT testing, the mice received a learning trial to freely explore two matching acrylic 4cm cubes that were either both smooth or both textured. Following a 5-minute inter-trial interval (ITI) in which the animals were removed to holding cages, the mice received a 3-minute test trial during which one of the cubes was replaced with a novel cube identical in appearance to the original object but with different tactile properties (smooth vs. textured). NOR was conducted the same as NORT except the objects differed visually, tactilely, and in size and materials, and the ITI was 50 minutes. The objects consisted of a ½ inch diameter white PVC standing pipe measuring 14 cm tall surrounded by a metal spiral and a 3D-printed blue block measuring 14.4 cm x 5 cm x 2.5 cm. For both NORT and NOR, object type and side on which the novel object was presented was counterbalanced across groups. The movement of the mice was tracked with ANY-maze Software (Stoelting, Co.). The outcomes analyzed included total distance traveled and time spent investigating the objects, defined as the nose within 10 mm zone surrounding the object and pointing towards the object, excluding any time the mouse was climbing on the object. All objects and the apparatus were cleaned with 0.02% chlorhexidine between trials.

##### One-hour locomotor activity

P900L (n=21, 11 male and 10 female) and litter-matched WT (n=21, 10 male and 11 female) mice were used for the remainder of behavioral tests, which were performed by the Intellectual and Developmental Disabilities Research Center Animal Behavior Subunit at Washington University in St. Louis. Locomotor activity was measured in a transparent polystyrene enclosure (47.6 cm x 25.4 cm x 20.6 cm) by measuring photobeam breaks, as previously described (Maloney, Yuede, et al., 2019). Total ambulatory movement, vertical rearing behavior, and time spent in a 33 cm x 11 cm central zone were measured.

##### Sensorimotor battery

Walking initiation, balance (ledge and platform tests), volitional movement (pole and inclined screens), and strength (inverted screen) were measured as described previously (Chen et al., 2021). For the walking initiation test, mice were placed on the surface in the center of a 21 cm x 21 cm square marked with tape and the time for the mouse to leave the square was recorded. During the balance tests, the time the mouse remained on an elevated plexiglass ledge (0.75 cm wide) or small circular wooden platform (3.0 cm in diameter) was recorded. During the Pole test, mice were placed at the top of a vertical pole pointing upwards, and the time for the mouse to turn and descend the pole was recorded. During the inclined screen tests, the mouse was placed head-down on an elevated mesh grid, and the time to climb up the grid was recorded. During the inverted screen test, a mouse was placed on an elevated mesh grid, which was then inverted 180°, and the time to fall was measured. Tests lasted for 1 minute, except for the pole test which lasted 2 minutes. Data used for analysis are an average of two trials done on subsequent days.

##### Continuous and accelerating rotarod

Balance and coordination were assessed using the rotarod test (Rotamex-5, Columbus Instruments, Columbus, OH) as previously described (Maloney, Yuede, et al., 2019), using both constant rotation (5 rpm, 60 second maximum) and acceleration rotation (5-20 rpm, 180 second maximum) trials. Three sessions of testing consisting of two trials each were conducted, and trials were averaged. To focus the task on coordination rather than learning, testing sessions were separated by 4 days.

##### Morris water maze

To assess spatial learning, we performed the Morris Water Maze, consisting of cued trials, place trials, and probe trials as previously described (Maloney, Yuede, et al., 2019). Animals were placed in a large water-filled pool, and time and distance to reach an escape platform were measured (ANY-maze, Stoelting). Maximum trial duration was 1 minute. During cued trials, there was a visible escape platform that was moved to new locations for each trial, and the mice experienced 4 trials per day (separated by 30-minute inter-trial-intervals) across 2 days. Performance was analyzed in 2-trial blocks, with trials averaged. Three days later, animals were tested in place trials in which the escape platform was submerged in a consistent location, and there were numerous distal visual cues available. Place trials occurred daily for 5 days, consisting of 2 blocks of 2 consecutive trials. Trials within blocks were separated by a 30- second interval, and blocks were separated by 2 hours. Mice were released in different areas of the maze and required to use visual cues to find the hidden platform. Trial data were averaged across the trials within each day. One hour after the final place trial occurred, the probe trial took place, in which the platform was removed entirely. The mouse was released from the quadrant opposite to the learned platform location and allowed to swim in the task for one minute. Time spent in each quadrant, and the number of crossings over the zone the platform was previously in were recorded.

##### Elevated plus maze

Elevated plus maze tests were done as previously described (Maloney, Rieger, et al., 2019). In brief, the elevated apparatus contains a central platform (5.5 cm x 5.5 cm) with four arms extending from the central platform (each 36 cm x 6.1 cm). Two opposing arms were open and two have 15 cm tall opaque Plexiglas walls. Test sessions were conducted in a dimly lit environment with in which the mouse was able to freely explore the apparatus for 5 minutes. Position was measured with beam-breaks and time, distance, and entries into each zone were recorded and analyzed (MotoMonitor, Kinder Scientific).

##### Conditioned fear

Fear conditioning was performed as previously described (Maloney, Rieger, et al., 2019). Briefly, mice were habituated to an acrylic chamber (26 cm x 18 cm x 18 cm) that contained a metal grid floor, a LED light which remained on during trials, and a chamber odorant. During the training day, baseline measurements of freezing behavior were collected for 2 minutes. Then, once per minute, three training rounds occurred in which a 20-second 80 dB tone sounded for 20 seconds. During the last 2 seconds of the tone (conditioned stimulus) a 1.0 mA foot-shock (unconditioned stimulus) occurred. The next day, contextual fear was tested, in which the animals were placed in the same chamber with the same odorant with the testing light illuminated but no tones or shocks delivered. The following day, cued fear was tested, in which the animals were placed in a new opaque box with a new odorant. After a 2-minute baseline period with no tone, the same 80 dB tone was played for the remainder of the 8-minute trial. During all trials, freezing behavior were recorded and analyzed.

##### Statistical analysis for behavioral tests

Behavioral data were analyzed and plotted using GraphPad Prism 9.4.1. No consistent genotype by sex interaction effects were observed for any behavioral tests, therefore data were collapsed across sex. Statistical testing was performed using planned assay-specific methods, such as using unpaired Student’s T-Tests for single parameter comparisons between genotypes, and within-subjects two-way or three-way repeated-measures ANOVA for comparisons across timepoints. Individual timepoints within repeated measures tests were evaluated using Sidak’s multiple comparisons test. Unless otherwise noted, bar graphs and line graphs indicate mean ±SEM.

#### DNMT3A Protein and RNA Expression

Cortex tissue from P900L and WT animals (2 weeks old) were dissected in ice-cold PBS, flash frozen with liquid nitrogen, and stored at −80°C. Half of the cortex was used for protein expression measurement with western blotting, and the other half was used for RNA expression via RT-qPCR. Expression was assessed at 2 weeks of age because this is a timepoint with high postnatal expression.

##### Western Blotting

Western blotting was performed as previously described (Christian et al., 2020). WT and P900L (n=8/genotype, 4 males, 4 females) half-cortexes were homogenized with protease inhibitors (Buffer: 10mM HEPES pH 7.9, 10mM KCl, 1.5mM MgCl_2_, 1mM DTT, 10mM EDTA), and 1% SDS was added prior to boiling the samples for 10 minutes at 95°C. Subsequently, samples were spun at 15,000g for 10 minutes, and supernatant was run through a Wizard Column (Fisher, Wizard Minipreps Mini Columns, PRA7211), and protein concentration was measured using a Bradford assay. Samples were diluted in LDS sample buffer with 5% β-mercaptoethanol and boiled for 5 minutes before being run on a gel. An 8% acrylamide gel was used, and samples were run for 60 minutes at 125V before being transferred to a nitrocellulose membrane. Blots were blocked for 1 hour at room temperature in TBS-T with 3% bovine serum albumin, then immunostained with anti-DNMT3A (Abcam, 1:1000, ab13888) or anti-α-Tubulin (Abcam, 1:1000, ab52866) for 12-16 hours at 4°C. After washing membranes, they were incubated with secondary antibodies for 1 hour at room temperature in light-protected boxes (IRDye 800CW Goat anti-Rabbit, or IRDye 800CW Goat anti-Mouse, LI-COR Biosciences, 1:15,000, product numbers: 926-32211 and 926-32210 respectively). Primary and secondary antibodies were diluted in 3% Bovine Serum Albumin in TBS-T. Blots were imaged using the LiCOR Odyssey XCL system and quantified using Image Studio Lite software (LI-COR Biosciences). DNMT3A and α-Tubulin levels were normalized to a standard curve, and protein levels are expressed as normalized DNMT3A values divided by normalized α-Tubulin values to enable comparison of DNMT3A levels between blots. Significance was assessed using an unpaired Student’s T-Test.

##### qRT-PCR

RNA from WT and P900L (n=5/genotype, 3 males, 2 females) half-cortexes were isolated using the AllPrep DNA/RNA kits (QIAGEN, 80284), and RNAs were reverse transcribed using the using the High-Capacity cDNA Reverse Transcription Kit (Applied Biosystems). *DNMT3A* and *ACTB* were measured by qPCR using the Power SYBR™ Green PCR Master Mix and primers for *ACTB* (F:AAGGCCAACCGTGAAAAGAT, R:GTGGTACGACCAGAGGCATAC) or *DNMT3A* (F:GGCCTTCTCGACTCCAGATG, R:TTCCTCTTCTCAGCTGGCAC). The Ct of each primer set in each sample was calculated, and relative quantity was determined by comparing to a standard curve and then normalizing the *DNMT3A* signal to the *ACTB* signal.

#### Whole Genome Bisulfite sequencing

##### Global methylation across brain regions

300ng of DNA was isolated from brain tissue from 8-week animals (n=2/sex/genotype/mutation/region) using the AllPrep DNA/RNA kit (QIAGEN, 80284). DNA was then fragmented for 45 seconds with the Covaris S220 sonicator (10% Duty Factory, 175 Peak Incidence Power, 200 cycles per burst, milliTUBE 200µL AFA Fiber). To select for long DNA inserts, DNA was purified using 0.7 volumes of Agencourt Beads. A small amount of Lambda DNA was spiked in to allow for estimation of non-conversion rates. To prepare bisulfite DNA libraries, we used the Tecan Ovation Ultralow Methyl-Seq Kit (Tecan, 0335-32) and the Epitect Bisulfite Kit (Qiagen, 59824). Alternate bisulfite conversion cycling conditions were used to ensure lowest possible non-conversion rate ([95°C, 5 min; 60°C, 20 min] x 4 cycles, 20°C hold). Libraries were PCR amplified 11-13 cycles and pooled for low-depth sequencing at the Washington University in St. Louis Center for Genomic Science. Libraries were sequenced using a MiSeq 2x150 and sequenced at an average depth of 0.018x genomic coverage (average 0.2M reads per sample). Sequencing data were processed as described below, and genome-wide averages of mCA and mCG were analyzed using a paired Student’s T-Test with Bonferroni correction.

##### Deep sequencing of cortical DNA methylation

50ng of DNA isolated from a total of sixteen 8-week cortex samples (n=2/sex/genotype/mutation) and fragmented for 45 seconds using the Covaris E220 sonicator (10% Duty Factory, 175 Peak Incidence Power, 200 cycles per burst, milliTUBE 200µL AFA Fiber) and purified using 0.7 volumes of SPRISelect Beads (Beckman Coulter Life Sciences). A small amount of Lambda DNA was spiked in to allow for estimation of non-conversion rates. DNA was then bisulfite converted using the EZ DNA Methylation-Direct Kit (Zymo Research Corporation, D5020) using extended bisulfite conversion incubation to ensure lowest possible non-conversion rates (98°C, 8 min; 64°C, 4 hours 15 min). Samples were either stored overnight at −20°C, or libraries were immediately prepared using the Accel-NGS Methyl-Seq DNA Library Kit (Swift, 30024) with combinatorial dual indexes (Swift, 38096) as instructed, using 10 cycles of final amplification. Libraries were pooled and sequenced at the Genome Technology Access Center at the Washington University McDonnell Genome Institute using the NovaSeq 6000 2x150. An average sequencing depth of 10x genomic coverage (average 144M reads per sample) were obtained per sample.

##### Whole genome bisulfite analysis

Analysis of bisulfite sequencing was performed as described previously (Christian et al., 2020; Clemens et al., 2019). Reads were adapter-trimmed, mapped to mm9, deduplicated, and called for methylation using BS-seeker2 (W. Guo et al., 2013). Bedtools map -o sum was used to assess methylation across regions, summing the number of reads mapped to Cs (interpreted as mC after bisulfite conversion) and then dividing by the sum of Cs and Ts (indicating C) at that region. %mC values from biological replicates were averaged together. Though our methods should maximize the amount of efficient bisulfite conversion, a small percentage of unmethylated cytosines can are called as methylated due to nonconversion (0.2-0.3%). To adjust for nonconversion rate, regions were adjusted by the % methylation measured in Lambda spike-ins per sample, similar to previous analysis (Lister et al., 2013). If corrected region values were below 0, the %mC value was set to 0. Due to background nonconversion, lowly methylated regions (e.g., mCA at CpG islands or promoters) are not expected to show the same percentage reduction in methylation as higher mCA regions.

##### Differentially methylated region detection

We used BSmooth (Hansen et al., 2012) on four biological replicates of P900L or R878H and their sex-matched WT littermates to call differential CpG methylated regions. CG sites were then filtered, requiring >2x genomic coverage in all replicates. Differentially methylated regions (DMRs) were called using a statistical threshold of t-stat >2.0, requiring length >100 bp, and biological replicate consistency (i.e. for hypomethylated regions, all WT mCG/CG values must be higher than mutant mCG/mCG values). Data fit the assumptions and requirements for BSmooth and fisher’s exact testing. Resampling for overlap analysis was done using bedtools shuffle.

#### Chromatin immunoprecipitation sequencing

##### Chromatin immunoprecipitation library generation

Chromatin immunoprecipitation was performed as previously described (Clemens et al., 2019). Cerebral cortex was dissected on ice in PBS from DNMT3A mutants and their WT littermates at 8-weeks old (n=2/sex/genotype/mutation; a total of 4 WT and 4 mutants in P900L litters, and a total of 4 WT and 4 mutants in R878H litters). The tissue was flash-frozen in liquid nitrogen and stored at −80°C. Chromatin were fragmented with the Covaris E220 sonicator (5% Duty Factory, 140 Peak Incidence Power, 200 cycles per burst, milliTUBE 1mL AFA Fiber). ChIP was performed with H3K27ac antibody (0.1µg; Abcam, ab4729) and libraries were generated using Accel-NGS 2S Plus DNA Library Kit (Swift Biosciences). Pooled libraries were sequenced using a NovaSeq 6000 with the Genome Technology Access Center at Washington University in St. Louis, typically yielding 20-50 million (average: 34 million) single-end reads per sample.

##### Chromatin immunoprecipitation analysis

ChIP sequencing analysis was performed as previously described (Clemens et al., 2019). In brief, reads were mapped to mm9 with bowtie2, and deduplicated with picardtools MarkDuplicates. Bedtools coverage -counts was used to assess H3K27ac signal at the various genomic regions examined. edgeR was then used to determine differential H3K27ac signal between WT and mutant animals. Data were visualized using the UCSC genome browser (Haeussler et al., 2019).

#### RNA sequencing

##### RNA sequencing library generation

Total RNA isolation was carried out as previously described (Clemens et al., 2019). In brief, cerebral cortex was dissected in ice-cold PBS from P900L or R878H mutants and their respective WT littermates at 8 weeks of age (n=7 pairs, 3 male, 4 female). Cortex was lysed in RLT buffer and RNA was isolated using the AllPrep DNA/RNA kit (QIAGEN, 80284). RNA libraries were generated from 250ng of RNA with NEBNext Ultra Directional RNA Library Prep Kit for Illumina (NEB) using a modified amplification protocol (37°C, 15 minutes; 98°C, 30 seconds; [98°C, 10 seconds; 65°C, 30 seconds; 72°C, 30 seconds]x13; 72°C, 5 minutes; 4°C hold). RNA libraries were pooled at a final concentration of 10nM and sequenced using Illumina NextSeq-High 1x75bp with the Center for Genome Sciences at Washington University in St. Louis, typically yielding 15-30 million single-end reads per sample.

##### RNA sequencing analysis

RNA sequencing analysis was performed as previously described (Clemens et al., 2019). Briefly, raw FASTQ files were trimmed with Trim Galore and rRNA sequences were filtered and removed with Bowtie. Remaining reads were aligned to mm9 using STAR (Dobin et al., 2013), and uniquely mapping reads were converted to BED files and separated into intronic and exonic reads. These exonic BED files were used to assess gene counts using bedtools coverage - counts.

##### Differential gene expression

DESeq2 was used to identify differentially expressed genes between mutants and their WT littermates. To control for batch, sex, and litter, paired analysis was done using a design = ∼ pair + genotype, and contrasted by genotype for all analysis. Though all libraries were processed in groups that contained P900L and R878H pairs, P900L and R878H datasets were analyzed separately. Significantly dysregulated genes were called when *p_ad_*_j_<0.1. Mutant-specific genes were defined as significantly regulated in one direction in one mutant, and either being unchanged (nominal p-value > 0.5) or regulated in the opposite direction in the other mutant.

##### Defining shared TBRS genes

RNA-seq data in the DNMT3A KO/+ (n=7 pairs; 4 male, 3 female) from Christian et al., 2020 were combined with P900L (n=7 pairs; 4 male, 3 female) and R878H (n=7 pairs; 4 male, 3 female) datasets. All datasets were generated from 8-week cortex and processed using similar methods. Datasets were then combined, and littermate pairwise genotype comparisons were made using DESeq2 across all WT and mutant animals (design = ∼ pair + group and contrasted by group; group defined as WT or mutant with no indication of origin dataset).

##### PANTHER Gene Ontology analysis

Gene set enrichment analysis was done using the PANTHER Overrepresentation Test (Version 17.0, Released 2022-02-22). Analyzed lists (e.g., significantly upregulated genes in the P900L mutant) were compared to a reference list of all expressed genes in our study (defined as genes with more than an average of 10 counts in both WT littermate datasets). Analysis identified PANTHER GO-slim Biological Process terms and used a Fisher test with FDR correction. A subset of the most significant PANTHER terms is shown in figures with full PANTHER results in Supplemental Table 2.

##### Rank-rank hypergeometric overlap (RRHO) analysis

For each mutant-WT pair, a ranked gene list was created using a gene score calculated as - log_10_(p-value) * sign (log_2_Fold Change) using the DESeq2 results for that gene. RRHO2_initialize() was used to generate RRHO object, and RRHO2_heatmap() was used to generate a heatmap of overlapping genes between different mutants.

#### Experimental design

Sample sizes were chosen based upon previously published studies using similar techniques. Statistical tests and exclusion criteria (values beyond 2 standard deviations of the group mean) were similar to that of previously published studies and indicated in the appropriate methods. For all animal experiments, experimenters were blinded to genotype during data collection. No treatment conditions were used, so no samples or animals were allocated to experimental groups and no randomization was needed. Tests that assume equal variance were only run if group variances were similar, otherwise alternative tests were used.

#### Data availability statement

The data that support the findings of this study are available from the corresponding author upon request. DOIs for all published gene sets used in comparison and enrichment analysis:

Lister et al. 2013: https://doi.org/10.1126/science.1237905

Clemens et al. 2019: https://doi.org/10.1016/j.molcel.2019.10.033

Christian et al. 2020: https://doi.org/10.1016/j.celrep.2020.108416

Hamagami et al. 2023: https://doi.org/10.1101/2023.02.17.528965

Bisulfite-seq, RNA-seq, and ChIP-seq are available on the NCBI GEO archive GSE225372.

Reviewer token: wjgnmggwhvkrjkr.

## Acknowledgements

We thank T. Ley and the TBRS community for reagents and discussion. We also thank J. Hoisington-Lopez and M. Crosby at the Center for Genome Sciences, GTAC for sequencing support, D. Wozniak and the IDDRC@WUSTL Animal Behavior Subunit, C. Semenkovich and S. Adak at the Washington University Diabetes Research Center, M. Brodt and the Washington University Musculoskeletal Research Center, and M. Haywood for assistance landmarking skulls. We thank our funding sources: NIH-NICHD F31HD100098 (to D.C.B.); NIH-NICHD P50HD103525 (to the IDDRC@WUSTL); and The Simons Foundation Autism Research Initiative, NIH-NIMH R01MH117405, and NIH-NINDS R01NS04102 (to H.W.G.).

## Author Contributions

Conceptualization and Methodology, DCB and HWG; Formal Analysis, DCB, DYW, NH, XG, ABH, ABL, CAH, TP, AM, JG, JDD, SEM; Investigation, DCB, XZ, JRM, NH, RGS, KBM, XG, ABH, HZ, ABL, TP, SEM; Writing – Original Draft, DCB and HWG; Writing – Review & Editing, all authors.

## Supplemental Information

Supplemental Information includes three figures and two tables.

Table S1. Detailed Statistical Methods and Outputs; Related to Figures 1-5, 7, S1-S3

Table S2. Full table of significant PANTHER terms, Related to Figures 6, 7

---

**Supplemental Figure 1:**
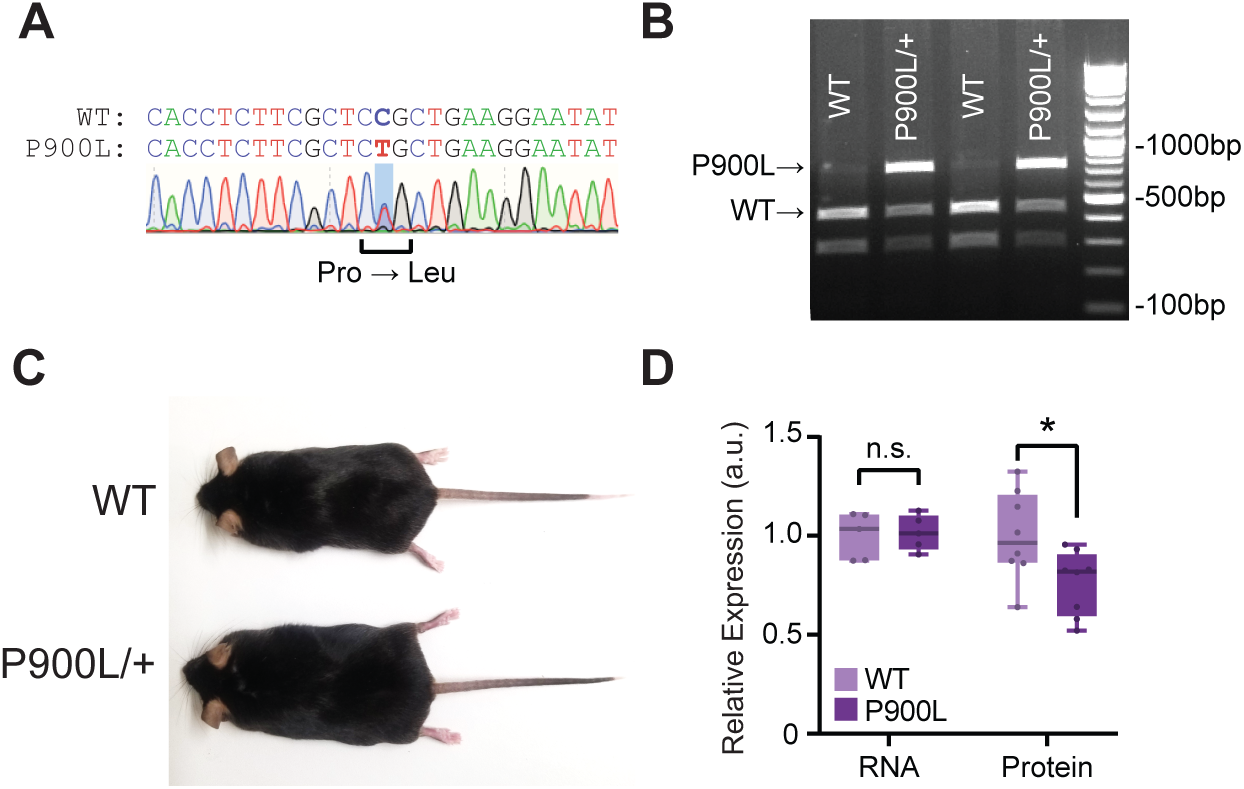
Generation of the P900L mutant model. (A) Sanger sequencing tracks indicating heterozygous P900L point mutation. The WT CCG codon is changed to a P900L CTG codon, with the mutation highlighted in blue. (B) Example gel from restriction enzyme genotyping from WT and P900L/+ ear lysate. Digested WT PCR products are at approximately 285bp and 421bp, whereas P900L PCR products are undigested and remain at the full 706bp. (C) Representative image of WT and P900L mouse at 30 weeks of age. (D) Quantification of DNMT3A expression from 2-week cortex by RT-qPCR (n=5/genotype, 3 males, 2 females) and western blot (n=8/genotype, 4 males, 4 females). Student’s T-Test; * p<0.05. Box plot indicates 25^th^ percentile, median, and 75^th^ percentile. Whiskers indicate minimum and maximum.

**Supplemental Figure 2:**
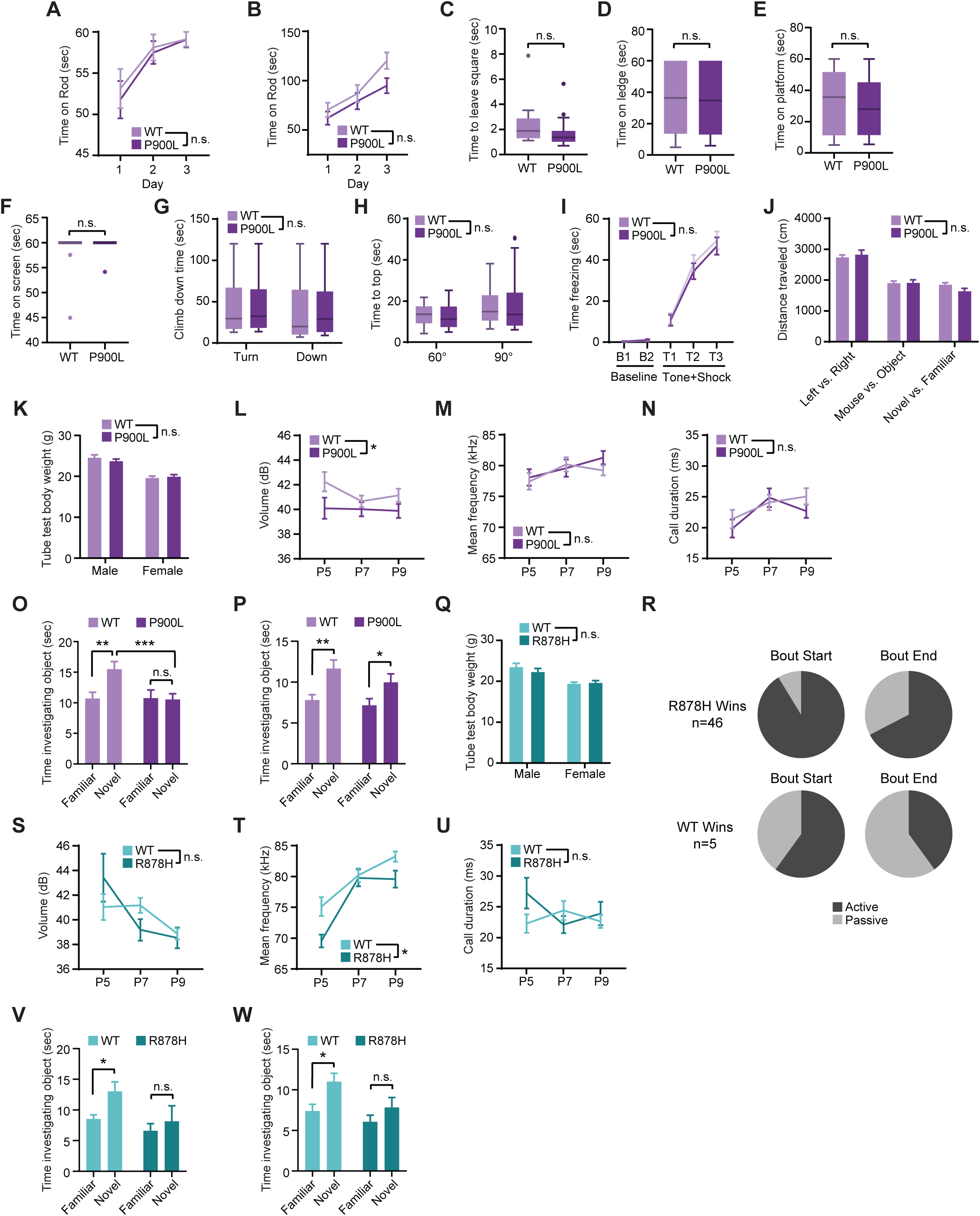
P900L mutants do not show activity or anxiety-like phenotypes, but do have changes in social and tactile behaviors. (A-B) Time on a continuous (A) or accelerating (B) rotarod indicates no significant differences between WT and P900L animals. (C) Walking initiation assay, with time to leave a marked square measured. (D-F) Genotypes had no significant differences in latency to fall of a (D) ledge, (E) platform, or (F) an inverted screen. (G-H) Genotypes had no significant differences in time taken to (G) turn and climb down a pole, or (H) time to the top of a 60° or 90° screen. (I) Time freezing during conditioned fear training, during baseline (before tone and shock) and during tone and shock association training. (J) Distance traveled during the 3-chamber social approach assay. (K) Body weights of animals during tube test assay indicate no significant differences between genotypes for the P900L animals vs. WT littermates. Body weight and size can have a significant impact on social hierarchies, and testing was done before mutants increased in size. (L-N) Measures of volume (L), average frequency (M), and duration (N) of ultrasonic vocalization calls in the WT and P900L animals. (O-P) Time spent investigating objects in NORT (O) and NOR (P) trials for WT and P900L animals. (Q) Body weight of R878H animals vs. WT littermates during tube test trials. (R) Active vs. Passive animal status during R878H or WT wins (see methods). (S-U) Measures of volume (S), average frequency (T), and duration (U) of ultrasonic vocalization calls in the WT and R878H animals. (V-W) Time spent investigating objects in NORT (V) and NOR (W) trials for WT and R878H animals. Bar graphs and line plots indicate mean ± SEM; Box-and-whisker plots indicate mean and quartiles. Detailed statistics, and sample sizes in Supplemental Table 1.

**Supplemental Figure 3:**
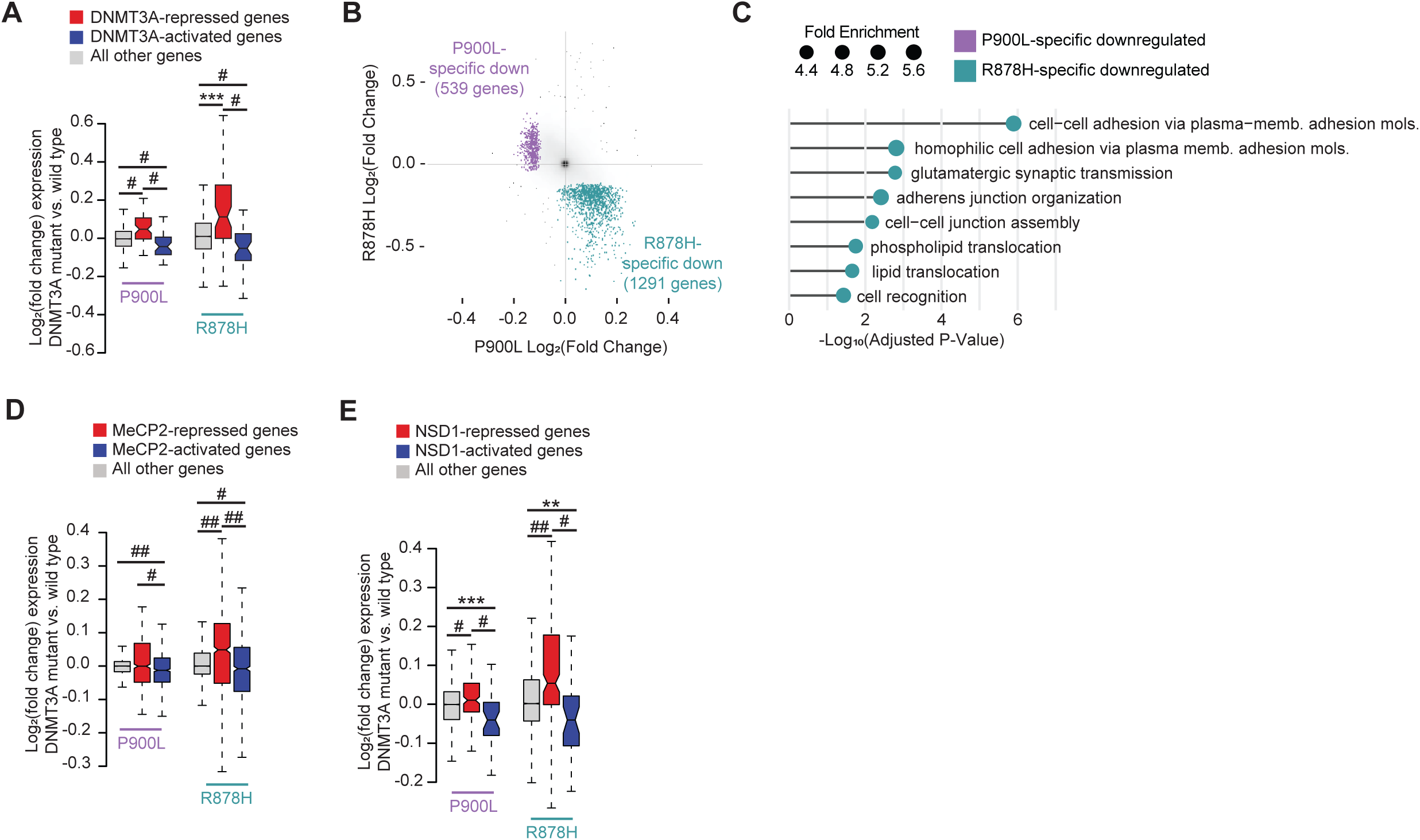
P900L and R878H mutants exhibit transcriptional overlap with other models disrupting the neuronal epigenome, and mutants also exhibit specific downregulated effects. (A) Log_2_ fold changes in the P900L- and R878H- mutants at genes significantly disrupted upon homozygous KO of DNMT3A in postmitotic neurons (Clemens *et al*., 2019) (B) P900L- and R878H-specific downregulated gene sets indicated in purple and teal. Specific genes are defined as those that are significantly downregulated in one mutant, and either significantly unchanged (nominal p-value > 0.5) or upregulated in the other (fold change > 0). (C) Most significant PANTHER gene ontology (biological process) terms enriched in P900L- specific and R878H-specific downregulated gene lists. No terms were significantly enriched in the P900L-specific downregulated gene sets. (D) Log_2_ fold changes in the P900L- and R878H- mutants at genes significantly disrupted in MeCP2 mutants (Clemens *et al*., 2019) (E) Log_2_ fold changes in the P900L- and R878H- mutants at genes significantly disrupted in upon homozygous KO of NSD1 in neural progenitors (Hamagami *et al*., 2023) Notched box and whisker plots indicate median, interquartile, and confidence interval of median with significance from Wilcox Rank Sum test shown. Detailed statistics, and sample sizes in Supplemental Table 1. ** *p*<0.01; *** *p*<0.001; # *p*<0.0001; ## *p*<2 e^-10^

